# Cardiac Mitochondrial Dysfunction Following Bleomycin-Induced Acute Lung Injury in Rats

**DOI:** 10.64898/2026.05.06.723353

**Authors:** Reesa M. Wilcox, Alice Ngu, Isabella A. Jiang, Grace K. Nielsen, Peter R. Pellegrino, Han-Jun Wang

## Abstract

**Background:** Acute lung injury (ALI) and acute respiratory distress syndrome (ARDS) are frequently associated with cardiac complications, including myocardial injury and right ventricular dysfunction. However, the mechanisms linking pulmonary injury to cardiac dysfunction remain incompletely understood. In this study, we investigated ventricular mitochondrial respiratory function during the acute phase of bleomycin-induced ALI.

**Methods:** ALI was induced in male and female rats by intratracheal bleomycin (2.5 mg/kg); saline served as a control. Circulating cardiac troponin I (cTnI) was measured as an indicator of myocardial injury. Mitochondrial respiration was assessed in permeabilized ventricular fibers using high-resolution respirometry (HRR). The mitochondrial respiration rate of the H9C2 cardiomyoblast cell line was performed using Seahorse X^fe^96 Cell Mitochondrial Stress Test. Cells were treated with pro-inflammatory cytokine cocktails (PRO; IL1β plus TNFα plus IL6), anti-inflammatory cytokine cocktails (ANTI; IL4 plus IL10), a mixture of PRO and ANTI (BOTH), and (-)-norepinephrine (NE) in either hypoxic (1% oxygen) or normoxic conditions.

**Results:** Bleomycin-induced ALI increased circulating cTnI levels in male rats, indicating early cardiac stress following lung injury. Mitochondrial respiration in the LV appeared to show modest alterations, with preserved oxidative phosphorylation (OXPHOS) and electron transport (ET) capacity. In contrast, the RV of male animals demonstrated marked reductions in absolute respiratory flux and substrate-supported OXPHOS capacity, indicating impaired mitochondrial oxidative capacity. Female animals exhibited greater preservation of mitochondrial respiratory function, particularly in the RV, with higher OXPHOS capacity and greater Complex I gain than males. H9C2 cells treated with PRO showed a significant increase in uncoupled respiration following 6- and 24-hour incubation periods, under normoxic conditions. Maximal respiration and spare respiratory capacity were increased following 24-hours under hypoxia. No significant changes were observed following treatment with NE alone and in combination with PRO under normoxia or hypoxia for 24 hours.

**Conclusions:** ALI induces ventricle-specific and sex-dependent alterations in cardiac mitochondrial bioenergetics, with pronounced impairment in males and relative mitochondrial resilience in females. In H9C2 cardiomyoblasts, short-term exposure (6–24 hours) to pro-inflammatory cytokines enhances uncoupled mitochondrial respiration under normoxic conditions, while short-term hypoxic exposure independently increases maximal respiration and spare respiratory capacity.

## Introduction

Acute respiratory distress syndrome (ARDS) is a severe form of acute lung injury (ALI) that affects approximately 3 million people each year, contributing to 10% to 15% of intensive care unit (ICU) admissions^1^. This potentially life-threatening condition affects roughly 24% of mechanically ventilated patients. Mortality rates remain high, between 35% and 46%, depending on severity at the time of presentation^2^. The systemic effects of ARDS and ALI, particularly through heart-lung interactions, have gained increasing recognition.

Cardiac complications such as myocardial damage, systemic inflammation, hypoxemia, and elevated pulmonary artery pressure that arise from severe ARDS are associated with worse clinical outcomes^3,4^. Right ventricular (RV) dysfunction, often precipitated by elevated pulmonary vascular resistance, greatly contributes to mortality^1,5^. Left ventricular (LV) diastolic dysfunction is associated with greater lung edema and contributes to lowered cardiac output and systemic hypoperfusion^6^.

The COVID-19 pandemic has more distinctly revealed the vulnerability of cardiac function during pulmonary distress. Approximately one third of COVID-19 patients admitted to the ICU developed biventricular dysfunction^7^. Despite increasing recognition of the cardiac sequelae of ALI, major gaps remain in our understanding of how ALI and ARDS affect the heart at a mechanistic level. The differential effects of pulmonary insults on the left and right ventricles have received limited attention, despite established differences in their structural, metabolic, and hemodynamic characteristics. Relatively few studies have addressed alterations in cardiac mitochondrial function following ALI, and fewer still have evaluated whether these changes differ between ventricular compartments. Experimental models of ALI, including those induced by bleomycin, acid aspiration, or endotoxin exposure, have primarily focused on pulmonary outcomes, limiting characterization of systemic bioenergetic consequences, particularly in the myocardium. However, some evidence suggests that pulmonary inflammation can impair cardiac mitochondrial respiration and increase oxidative stress^8^. The broader metabolic remodeling of the heart in response to lung injury is insufficiently studied. These outcomes reveal the need to evaluate cardiopulmonary interactions at the level of mitochondrial function and transcriptional remodeling.

In our previous work, we demonstrated that bleomycin-induced ALI elicits spontaneous arrhythmias, particularly premature ventricular contractions (PVCs), emerging in one week after bleomycin injury, peaking by week 3, and resolving by week 4^9^. Although this study suggested that neural inflammation in the stellate ganglia mediates PVC susceptibility during ALI recovery, the molecular and cellular mechanisms underlying spontaneous PVC generation post-ALI remain unclear. In the present study, we focus on the heart itself as a site of secondary injury to address the mechanism of PVC development. We hypothesize that bleomycin-induced ALI causes direct cardiomyocyte stress and mitochondrial dysfunction during the early post-injury period, contributing to electrical instability and metabolic vulnerability. Myocardial bioenergetics were assessed using Oroboros O2k high-resolution respirometry (HRR). The potential effects of hypoxia, inflammatory cytokines, and sympathetic excitation-released neuronal transmitter norepinephrine (NE) on cardiac mitochondrial function were tested in H9C2 cardiomyocytes.

Recent studies indicate that sex-based differences in mitochondrial function may modulate susceptibility to cardiac injury during critical illness. Clinically, female patients have demonstrated a reduced incidence and severity of ARDS, which may be attributed to hormonal factors and mitochondrial protective mechanisms^10,11^. Preclinical work has demonstrated sexual dimorphism and the effects of sex hormones on mitochondrial regulation in cardiac tissue, including notable patterns in the expression of genes encoding mitochondrial proteins, reactive oxygen species (ROS) buffering, and respiratory capacity^12,13^. The second goal of this study is to examine the potential sex difference in cardiac mitochondrial stress post-ALI.

## Methods

### Ethical Approval

Animals were housed in a temperature-controlled environment (22°C–25°C) with a 12-hour light-dark cycle and provided ad libitum access to food, water, and environmental enrichment. All experimental protocols were approved by the Institutional Animal Care and Use Committee (IACUC) of the University of Nebraska Medical Center (protocol ID no. 23-012-03 FC) and were in accordance with the National Institutes of Health guidelines for the Care and Use of Laboratory Animals.

### Animal Husbandry and Housing

Sprague-Dawley rats (2–3 months old) were used in all experiments. Animals were housed in groups on a 12-hour light/dark cycle, with access to food, water, and enrichment materials, and were monitored regularly for health and welfare. All experiments were conducted during the light phase (09:00–16:00 h).

### Intratracheal Instillation of Bleomycin

All rats underwent intratracheal treatment of bleomycin (Bleo) or a saline vehicle (Sham). Rats were anesthetized using 2%–3% isoflurane in oxygen. An endotracheal tube was orally inserted, and bleomycin sulfate (2.5 mg/kg body weight) was administered intratracheally in 150 µl of sterile saline. Control animals received an equal volume of intratracheal sterile saline.

### Animal Cohorts

Two independent animal cohorts were utilized in this study. The first cohort was used to measure plasma cardiac troponin I (cTnI) levels and included male Bleo (n = 9), male Sham (n = 8), female Bleo (n = 11), and female Sham (n = 8). The second cohort was used for HRR and comprised 28 rats, equally divided by sex: male Bleo (n = 8), male Sham (n = 6), female Bleo (n = 7), and female Sham (n = 7). Cardiac tissue for this cohort was collected 7–10 days post-treatment.

### Cardiac Troponin I Measurement

Rats in the first cohort underwent cTnI measurement. The artery on the ventral aspect of the rat tail was used to collect small amounts of blood (∼0.1 mL) for arterial blood gas analysis at day 7 post-Bleo or Sham. The animal was restrained with a commercial restrainer so that its tail was accessible. The tail was prepared aseptically by alternating alcohol prep pads and iodine prep pads three times, and the artery was then punctured using a 24 G needle. A small volume of blood (∼0.1 mL) was gently aspirated into the syringe for blood gas analysis (iSTAT, Abbott, Chicago, IL, USA). After sample collection, the needle was removed, and pressure was applied with sterile gauze to achieve hemostasis at the puncture site.

### Permeabilized Muscle Fibers

Rats in the second cohort were anesthetized 7-10 days after intratracheal treatment with 2%–3% inhaled isoflurane until unconsciousness was achieved. Then, an intraperitoneal injection of a euthanasia solution containing Urethane (7 g) and chloralose (0.7 g) dissolved in 50 ml saline (2 cc per 5 kg body weight) was administered. The thoracic cavity was opened, and the heart was rapidly excised, briefly weighed, and placed on ice. Tissue from the middle of the LV and RV were dissected and placed in ice-cold BIOPS buffer (10 mM Ca2+-EGTA–0.1µM free Ca2+, 20mM imidazole, 20 mM taurine, 50 mM K+-MES, 0.5 mM dithiothreitol, 6.56 mM MgCl2, 5.77 mM ATP, 15 mM phosphocreatine, pH 7.1 adjusted with KOH) as previously described^14^. A standardized mechanical and chemical permeabilization protocol using 50 µg/ml saponin (Sigma-Aldrich) was used to prepare permeabilized muscle fibers (pfi) (Pesta & Gnaiger, 2012).

### High-Resolution Respirometry

HRR was performed on chemically cardiac pfi isolated from LV and RV 7–10 days following Bleo or Sham treatment. Permeabilization was performed as previously described after which the permeabilized cardiac muscle fibers were washed for 10 minutes in 4 mL of mitochondrial respiration buffer respiration medium ((Oroboros MiR05-Kit): 110 mM sucrose, 60 mM K+-lactobionate, 0.5 mM EGTA, 3 mM MgCl2, 20 mM taurine, 10 mM KH2PO4, 20 mM HEPES + 280 units ml-1 catalase (MiR06)) at 4°C on a rocker.

Oxygen consumption rates were measured using an Oroboros O2k polarographic oxygen sensor (Oroboros Instruments, Innsbruck, Austria), with 1-2 mg of permeabilized cardiac muscle fibers from each ventricle separately loaded into 2 mL MiR06 buffer in chambers A and B. The chambers were maintained at 37°C with continuous stirring at 750 rpm. Initial oxygen concentrations were raised to 400–500 µM by injecting pure oxygen into the gas phase using the Oxia device (Oroboros Instruments, Innsbruck, Austria).

Mitochondrial respiratory capacity was assessed using the substrate-uncoupler-inhibitor titration (SUIT) 008 O2 pfi D014 protocol to evaluate the additivity of NADH-linked (N-pathway) and succinate-linked (S-pathway) respiration at the Q-junction, with an extension to examine Complex IV (CIV) activity in ventricular tissue. Sequential titrations of substrates and inhibitors (Sigma Aldrich) were added as follows: 5 mM pyruvate, 2 mM malate, to measure the resting electrochemical proton gradient (Leak), followed by 7.5 mM ADP, 10 µM cytochrome c, 10 mM glutamate, and 10 mM succinate to stimulate oxidative phosphorylation (OXPHOS) capacity. Maximal electron transfer (ET) system capacity was induced via stepwise additions of 0.25 µM CCCP. Inhibition of mitochondrial complexes was performed by sequential addition of 0.5 µM rotenone (Complex I) and 2.5 µM antimycin A (Complex III). Stimulation of CIV was completed by 2.5 mM TMPD in ascorbate, and sequential inhibition by titration of 200 mM sodium azide.

Outer mitochondrial membrane integrity was validated by confirming that the increase in oxygen consumption following cytochrome c addition did not exceed 10%. Oxygen concentration was maintained between 200–400 µM throughout the protocol by periodic titration of 200 mM hydrogen peroxide (H₂O₂) as needed.

HRR was used to evaluate oxygen flux across several coupling control and mitochondrial pathway states corresponding to the SUIT-008 protocol. Leak (*L*) respiration is reliant on pyruvate + malate (PM*_L_*). OXPHOS (*P*) respiration is sustained by PM + ADP (PM*_P_*), PM + glutamate (PGM*_P_*), and PGM + succinate (PGMS*_P_*). ET (*E*) is supported by uncoupled respiration (PGMS*_E_*) and Complex II (S*_E_*; S-pathway). CIV respiration was determined by subtracting the respiration of inhibition from stimulation.

Derived metrics quantifying mitochondrial respiratory capacity, control, substrate support, and efficiency were calculated from baseline-corrected absolute oxygen flux values. To maintain data quality, individual respiration profiles were evaluated using a regression-based diagnostic comparing each trace to its group-specific mean vector. Profiles with extreme slopes, indicative of global scaling artifacts consistent with calibration errors, were excluded to ensure the final dataset represented physiological variation rather than technical bias.

### Immunoblot Analysis

Samples of LV and RV were taken from Sprague-Dawley rats. While on ice, 50-70 mg of tissue were weighed out for protein analysis and homogenized as whole tissue. Tissues were processed using Dounce glass tissue grinder (SP Wilmad-LabGlass, Vineland, NJ, USA) in 300 µL Pierce RIPA lysis buffer (Thermo Scientific, Waltham, MA, USA) and 3 µL protease inhibitor cocktail (Sigma-Aldrich, St. Louis, MO, USA). Following manual grinding, tissues were briefly sonicated three times for 5 seconds to form tissue lysates. Subsequently, lysates were centrifuged at 12,000 rpm for 20 min at 4 °C to precipitate debris, then stored in a new tube at −80 °C. Protein concentration of each sample was determined using the Pierce BCA Protein Assay (Thermo Scientific, Waltham, MA, USA). An equal amount of protein from each sample was subjected to SDS-polyacrylamide gel electrophoresis (SDS-PAGE) and transferred to polyvinylidene difluoride (PVDF) membranes. Total protein content was determined using Ponceau S Solution (Sigma-Aldrich, St. Louis, MO, USA). Proteins of interest were visualized using the iBright CL750 Imaging System (Thermo Fisher Scientific, Waltham, MA, USA) and immunodetected using SuperSignal West Femto Maximum Sensitivity Substrate (Thermo Scientific, Waltham, MA, USA). Bands of interest were then analyzed using Image Studio software (LI-COR Biosciences, Lincoln, NE, USA). The following antibodies and respective dilutions were used: Total OXPHOS Rodent WB Antibody Cocktail (ab110413, 1:10000) included NADH: Ubiquinone Oxidoreductase Subunit B8 (NDUFB8; CI), Succinate Dehydrogenase Complex Iron Sulfur Subunit B (SDHB; CII), Ubiquinol-Cytochrome C Reductase Core Protein 2 (UQCRC2; CIII), Mitochondrially Encoded Cytochrome C Oxidase I (MTCO1; CIV), or ATP Synthase F1 Subunit Alpha (ATP5A; CV), NADH: Ubiquinone Oxidoreductase Core Subunit S1 (NDUFS1; ab96428, 1:1000), Succinate Dehydrogenase Complex Flavoprotein Subunit A (SDHA; ab96428, 1:1000), Ubiquinol-Cytochrome C Reductase, Rieske Iron-Sulfur Polypeptide 1 (UQCRFS1; ab14746, 1:1000), BCL2 Interacting Protein 3 (BNIP3; ab109362, 1:1000), Voltage Dependent Anion Channels 1 and 3 (VDAC1 + VDAC3; ab14734, 1:1000) from Abcam (Cambridge, MA), and GAPDH (sc-32233, 1:5000) from Santa Cruz (Santa Cruz, CA). Results for protein expression were normalized to VDAC1 + VDAC3 (mitochondrial loading control).

### Cell culture and Treatments

The H9C2 cardiomyoblast cell line was cultured in complete medium consisting of DMEM (ATCC 30-2002, VA, USA) supplemented with 10% FBS (ATCC 30-2020, VA, USA) at 37°C in 5% CO_2_. The cells were subcultured before reaching confluence to prevent loss of the myoblast population. A density of 6.7 x 10^3^ H9C2 cells was seeded in Seahorse X^fe^96 cell culture microplates of Seahorse FluxPaks (Agilent 103793-100, CA, USA) and incubated for 48 hours at 37°C in the atmospheric O_2_ level and 5% CO_2_ before treatments. The cells were treated with pro-inflammatory cytokine cocktails (PRO) at a final concentration of 100 ng/ml each of interleukin (IL)-6 (R&D Systems 506-RL-050/CF, MN, USA), IL-1β (R&D Systems 501-RL-050/CF, MN, USA), and tumor necrosis factor (TNF)-α (R&D Systems 510-RT-050/CF, MN, USA), anti-inflammatory cytokine cocktails (ANTI) at a final concentration of 100 ng/ml each of IL-10 (R&D Systems 522-RLB-025/CF, MN, USA), and IL-4 (R&D Systems 504-RL-025/CF, MN, USA), or a mixture of PRO and ANTI (BOTH)] for 6 or 24 hours before measurement of mitochondrial respiration rate. The cells were also treated with a final concentration of 50 µM (-)-NE (Sigma-Aldrich A7257, MO, USA). For hypoxia, the seeded cells were incubated at 1% O_2_ in a hypoxia chamber, whereas incubation at atmospheric O_2_ was considered normoxia.

### Seahorse X^fe^96 Cell Mitochondrial Stress Test

The measurement of mitochondrial respiration rate was performed using the Seahorse X^fe^96 flux analyzer. After cell culture treatment(s), the cells were washed once and then incubated with pH7.4-adjusted XF base medium (Agilent 103335-100, CA, USA) supplemented with a final concentration of 10 mM glucose (Agilent 103577-100, CA, USA), 1 mM pyruvate (Agilent 103578-100, CA, USA), and 2 mM L-glutamine (Agilent 103579-100, CA, USA) in the incubator without CO_2_ for an hour. The mitochondrial stress test was performed following the default protocol commands for calibration and equilibration, and the default measurement cycle times: 3 cycles of 3 minutes mix, 0 minutes wait, 3 minutes measure for baseline measurement cycle, and each acute compound injection—oligomycin, FCCP, and Rotenone/Antimycin A (Agilent 103015-100, CA, USA) with the final concentrations of 1 µM, 4 µM, and 0.5 µM respectively. Cell lysate from each well was prepared, and protein concentration was measured using the BCA protein assay (Thermo Fisher Scientific 23225, MA, USA). Cell lysate mass-normalized oxygen consumption rate (OCR) was measured in each well. Parameters such as basal respiration, non-mitochondrial oxygen consumption, proton leak, ATP production, maximal respiration, and spare respiratory capacity were calculated.

### Statistical Analysis

Pathway-level Mitopathways analyses were conducted separately for LV and RV. Baseline-corrected flux values at each respiratory state were analyzed independently using two-way ANOVA with treatment and sex as fixed factors. Post-hoc significance testing within each state was performed using Šídák’s method for multiple comparisons to control the familywise error rate. Mixed-effects models with restricted maximum likelihood (REML) were applied when group sizes were unequal or data were excluded. Functional mitochondrial metrics, including gain, coupling, uncoupling penalty, substrate-supported ET, and control ratios, were analyzed using three-way ANOVA with treatment, sex, and ventricle as fixed factors, with ventricle treated as a matched (repeated-measures) factor to account for paired left and right ventricular measurements from the same animal. Mixed-effects models with REML were used as appropriate, and Šídák’s post-hoc testing was applied following significant main effects or interactions. Circulating cTnI concentrations were assessed using linear mixed-effects models with treatment and sex as fixed factors. Western Immunoblotting was analyzed using Mann-Whitney U test. Data are expressed as mean ± SEM; statistical significance was defined as *P* < 0.05.

Seahorse X^fe^96 Cell Mitochondrial Stress Test OCR parameters of basal respiration, non-mitochondrial oxygen consumption, proton leak, ATP production, maximal respiration, and spare respiratory capacity were quantified in H9C2, *Rattus Norvegicus* cardiomyoblast cell line. Two-way ANOVA was employed to analyzed main interaction and effects of the fixed factors oxygen level and chemical mediator treatment (NE and/or PRO treatment), followed by Tukey’s post hoc test. A similar statistical test was conducted on the 24-hour incubation of inflammatory cytokine cocktails with PRO and ANTI as fixed factors. Multiple comparisons between groups in *in vitro* experiments, such as a 6-hour inflammatory cytokine incubation experiment and a chemical mediator treatment experiment, were analyzed using one-way ANOVA with Dunnett’s multiple comparisons test relative to control (CTRL). Normoxia and hypoxia conditions were compared using unpaired two-tailed Student’s *t*-tests. Data are reported as mean ± SD. Statistical significance was defined as *P* < 0.05.

## Results

### Bleomycin-Induced ALI Elevates Circulating cTnI in a Sex-Specific Manner

cTnI concentrations were measured seven days post-Bleo to assess acute myocardial injury (**Figure 1**). In male rats, exposure to Bleo resulted in a significant increase in cTnI compared with Sham (*P* = 0.017), whereas in female rats, there was no difference between groups (*P* = 0.846). Sex comparisons showed that cTnI concentrations in Bleo males were significantly higher than Bleo females (*P* = 0.017).

**Figure 1.**
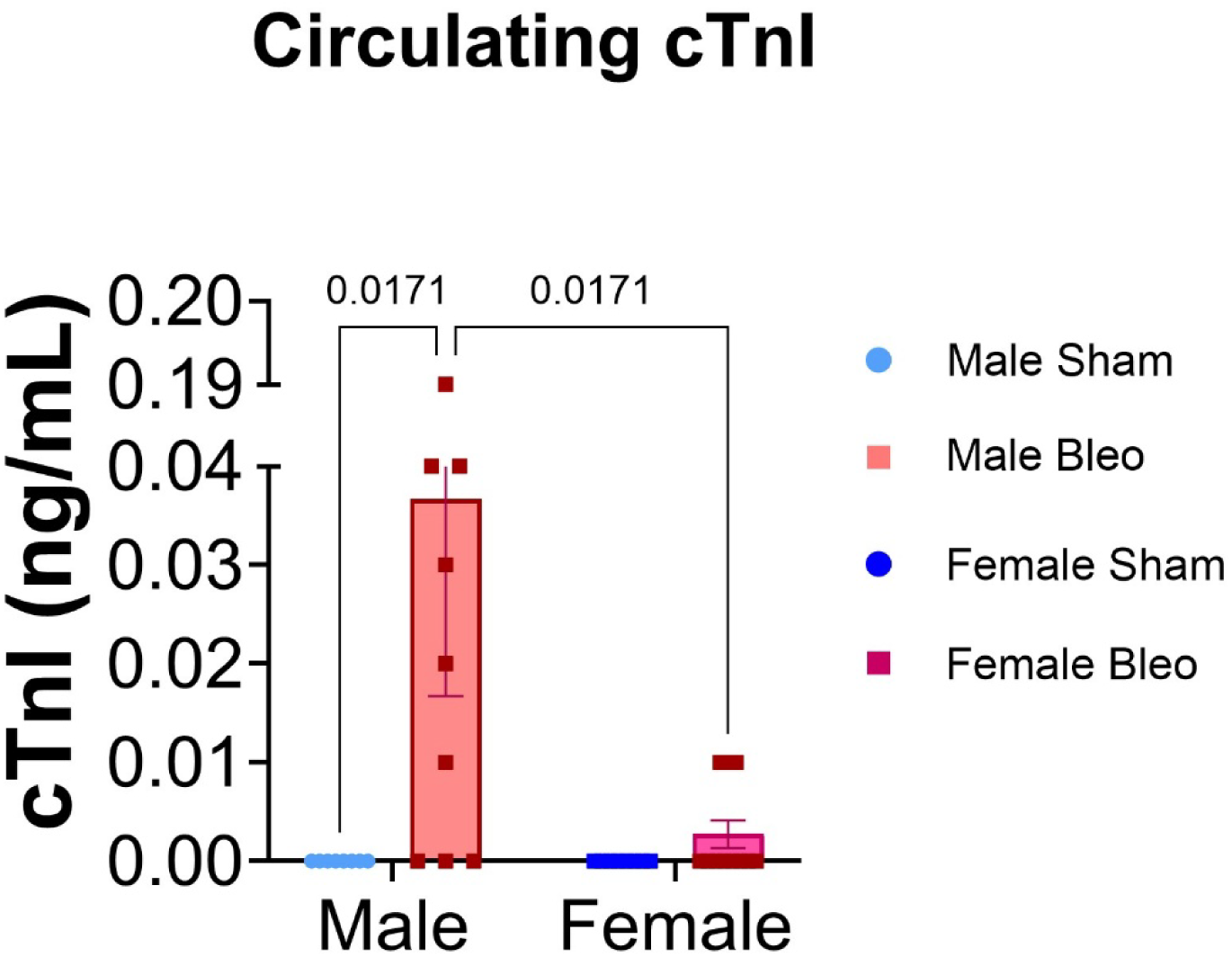
cTnI concentrations measured seven days post-Bleo resulted in significantly increased levels in male rats (n = 9) compared to Sham males (n = 8) and Bleo females (n = 11). No significance was observed between Bleo females and Sham females (n = 8). Data are expressed as mean ± SEM. Statistical analyses were performed using two-way ANOVA with sex and treatment as factors.

### Mitochondrial Respiratory Dysfunction in Left and Right Ventricles Post-ALI

Mitochondrial respiratory function was assessed in both left and right ventricular tissue using HRR 7-10 days after ALI was induced (**Figure 2**). Oxygen consumption values were normalized to wet tissue mass to allow for comparison of mitochondrial respiration across different substrates and respiratory states (**2A-C**). The LV of male Bleo rats showed reduced PGMS*_P_* (*P* = 0.029; **Figure 2B**) compared with Sham males. In the RV, Bleo males exhibited reduced PGMS*_P_* (*P* = 0.008) and marginal, non-significant reductions in PGMS*_E_* (*P* = 0.071) and S*_E_* (*P* = 0.089) compared to Sham males. When compared to Bleo females, Bleo males also exhibited significantly reduced PM*_L_* (*P* = 0.007) in the LV, PGM*_P_* (*P* = 0.021), PGMS*_P_* (*P* = 0.0097), and S*_E_* respiration (*P* = 0.036) in the RV. PM*_L_* trended lower in Bleo males compares to females (*P* = 0.077) but did not reach significance (**Figure 2C**).

**Figure 2.**
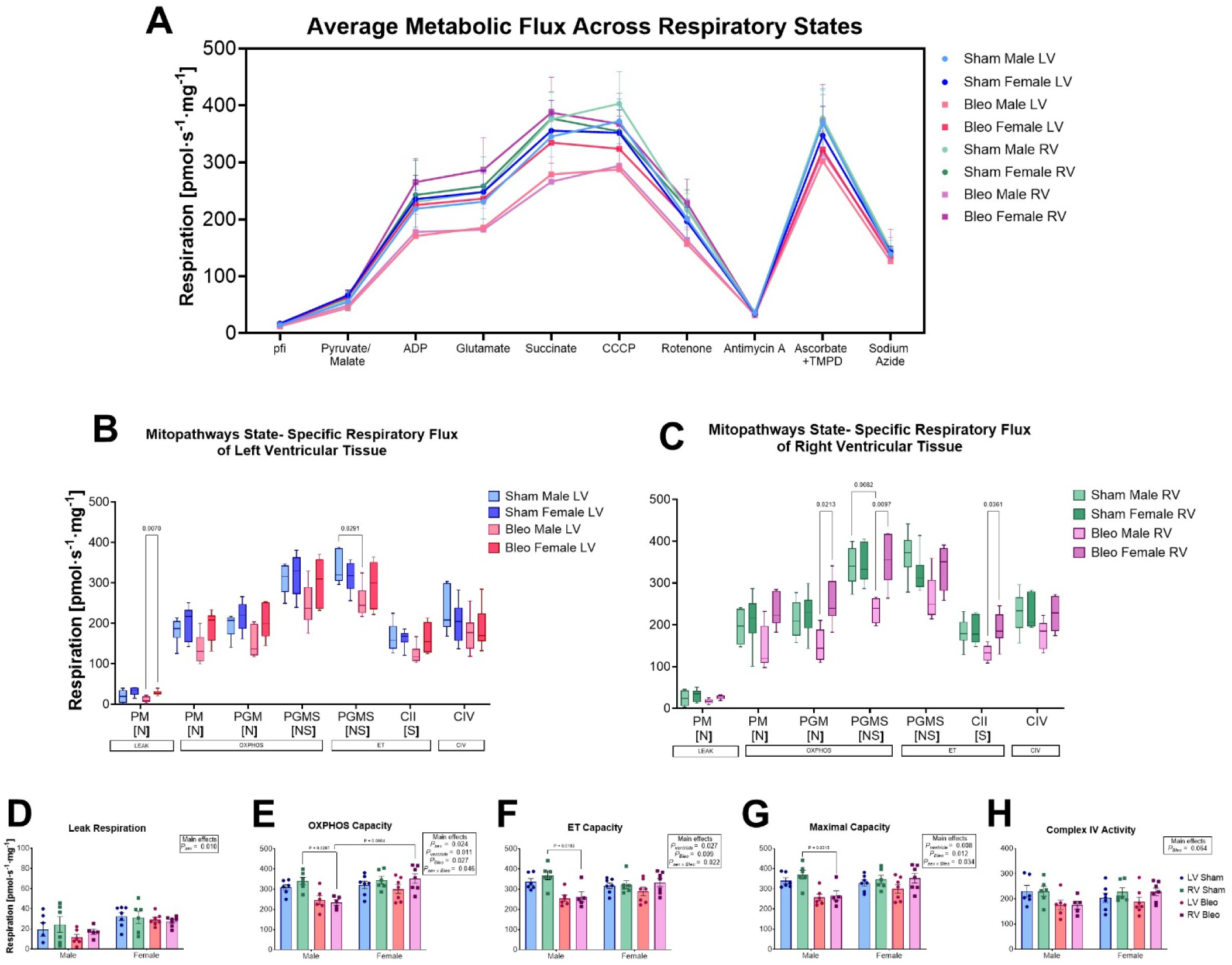
Average specific flux traces per group, measured throughout the experimental protocol and across the respiratory states of each combination of substrates, uncoupler, and inhibitors, are shown in (**A**). Mitopathways specific respiratory states leak (Leak; PM), oxidative phosphorylation (OXPHOS; PM, PGM, PGMS), electron transfer (ET; PGMS, CII), and complex IV activity (CIV) assessed in LV and RV tissue from rats with Bleo or Sham are shown in (**B**) and (**C**), respectively. Additivity between the N- and S-pathways, as denoted below, provides an estimate of maximum respiratory capacity. Quantitative comparisons of key respiratory metrics in (**D-H**) are baseline-corrected values. Maximal capacity indicates the maximal respiration value reached between succinate and CCCP (**G**). Complex IV activity was determined by subtracting the value of sodium azide from ascorbate + TMPD (**H**). Data are presented as mean ± SEM with individual data points (n = 5–7 per group). Statistical analysis for Mitopathways respiratory states in **(B)** and **(C)** was performed using two-way ANOVA with sex and treatment as factors, followed by Šídák’s multiple-comparisons test. Significance testing of respiratory metrics in **(D–H)** was completed using a three-way ANOVA with sex and treatment as between-subjects factors and ventricle laterality as a within-subjects factor followed by post-hoc testing using Šídák’s correction for multiple comparisons were analyzed using three-way ANOVA with sex, treatment, and ventricle as factors, with ventricle treated as a matched factor to account for paired LV and RV samples from the same animal, followed by Šídák’s multiple-comparisons test.

Residual oxygen consumption (ROX) baseline-corrected flux values were used to quantify absolute respiratory capacities across complexes and substrate combinations (**Figure 2D–H**). Three-way ANOVA revealed a significant main effect of sex on Leak respiration (*P_sex_* = 0.010; **Figure 2D**); however, post-hoc testing did not identify significant differences between groups. Significant main effects of sex (P_sex_ = 0.024), ventricle (*P_ventricle_* = 0.011), treatment (*P_Bleo_* = 0.027), and sex x treatment interaction (P_sex x Bleo_ = 0.046) were observed in OXPHOS capacity, as well as a reduced function in the RV of Bleo males compared to Sham males (*P* = 0.029; **Figure 2D**). ET capacity demonstrated significant main effects of ventricle (*P_ventricle_* = 0.027), treatment (*P_Bleo_* = 0.009), and sex x treatment interaction (P_sex x Bleo_ = 0.022), and reduced function in the RV of Bleo males compared to Sham (*P* = 0.018; **Figure 2E**). Congruently, maximal capacity demonstrated the same significant main effects (*P_ventricle_* = 0.008; *P_Bleo_* = 0.012; P_sex x Bleo_ = 0.034) and reduction of the RV in Bleo males compared to Sham (*P* = 0.032; **Figure 2G**).

Although neither remained significant following post-hoc testing, ET (*P* = 0.066; **Figure 2F**) and maximal capacity (*P* = 0.080; **Figure 2G**) trended lower in the LV of Bleo males compared to Sham. The RV of female Bleo rats trended an increase of OXPHOS (*P* = 0.058; **Figure 2E**) and maximal capacity (*P* = 0.077; **Figure 2G**) compared to LV but did not reach significance (*P* = 0.058; **Figure 2E**). A significant sex-specific difference in the RV was observed between Bleo males and females (*P* = 0.006; **Figure 2G**). CIV activity did not differ significantly between groups, although the treatment effect approached significance in the three-way ANOVA (*P_Bleo_* = 0.064; **Figure 2H**).

### Diminished Mitochondrial Respiratory Control and Complex I Deficits following ALI

We analyzed Complex I (CI)-linked respiratory metrics to assess early impairment in mitochondrial substrate utilization and NADH-driven respiration following ALI (**Figure 3**). OXPHOS gain, defined as ADP-stimulated respiration above Leak, showed a significant main effect of ventricle (*P_ventricle_* = 0.017) and a treatment effect approaching significance (*P_Bleo_* = 0.051) in the three-way ANOVA. Though no significant changes were identified in the LV or the RV of Bleo males, the RV OXPHOS gain in Bleo males trended lower compared to Sham males (*P* = 0.058). Additionally, the RV of Bleo females trended higher compared to the LV (*P* = 0.052); however, only the RV of Bleo males was significantly reduced compared with Bleo females (*P* = 0.019; **Figure 3A**).

**Figure 3.**
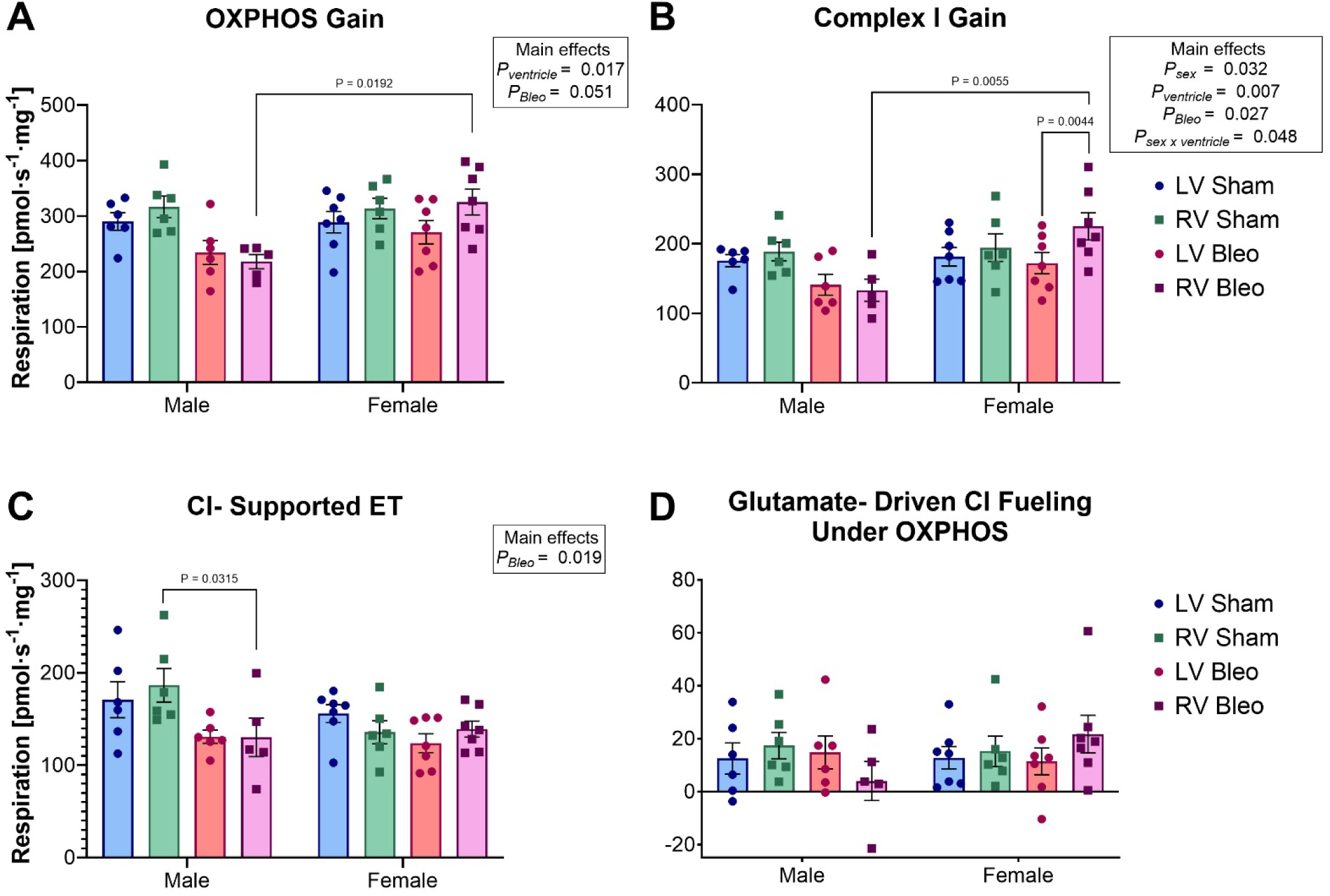
Complex I-linked respiration and substrate utilization in LV and RV tissue from male and female rats post-ALI are shown in (**A–D**). OXPHOS gain indicates ADP-stimulated respiration above Leak (**A**). CI gain denotes the contribution of NADH-linked substrates to OXPHOS capacity (**B**). CI-supported ET reflects uncoupled respiration supported by CI-linked substrates (**C**). Glutamate-driven CI fueling captures TCA cycle-linked support for CI-mediated respiration (**D**). Bleo males showed reduced RV OXPHOS Gain and CI Gain compared to Bleo females **(A, B),** and in CI-supported ET compared to Sham males (**C**). Bleo female rats exhibited reduced CI gain in RV compared to LV tissue (**B**). Data are expressed as mean ± SEM. Group sizes: n = 5–7 per experimental condition. Metrics were derived from baseline-corrected state-specific mitochondrial flux values. Significance testing was completed using a three-way ANOVA with sex and treatment as between-subjects factors and ventricle laterality as a within-subjects factor followed by post-hoc testing using Šídák’s correction for multiple comparisons.

CI gain, the NADH-dependent increase in OXPHOS capacity, showed significant main effects of sex (P_sex_ = 0.032), ventricle (*P_ventricle_* = 0.007), and sex x treatment interaction (P_sex x Bleo_ = 0.048). Consistent with no significant effect of treatment (*P_Bleo_* = 0.2694), no differences were observed between Bleo male and Sham male groups in either ventricle (*P* = 0.817 and *P* = 0.393, respectively). In Bleo females, RV CI gain was higher than LV (*P* = 0.004). Additionally, complex I gain was significantly higher in Bleo female RV compared to Bleo male RV (*P* = 0.006; **Figure 3B**).

CI-supported ET capacity, derived from uncoupled respiration with CI-linked substrates, showed a significant main effect of treatment in the ANOVA (*P_Bleo_* = 0.019). No significant changes were observed in the LV of Bleo males, but the RV was reduced compared with Sham males (*P* = 0.032). No significant differences were observed within the female groups (**Figure 3C**).

Glutamate-driven CI fueling, a marker of NADH production linked with the TCA cycle, trended lower in RV Bleo males but did not reach statistical significance compared with male Sham or Bleo females (*P* = 0.922 and *P* = 0.567; **Figure 3D**). These findings suggest selective impairment of CI-mediated respiratory control, particularly in the RV of male rats post-ALI.

### Reduced ET Reserve with Preserved Downstream Coupling following ALI

To determine if ALI affected later stages of mitochondrial respiration, we examined metrics of integrative control and downstream ET (**Figure 4**). Complex II (CII) recruitment by succinate under OXPHOS conditions did not differ across groups (**Figure 4A**).

**Figure 4.**
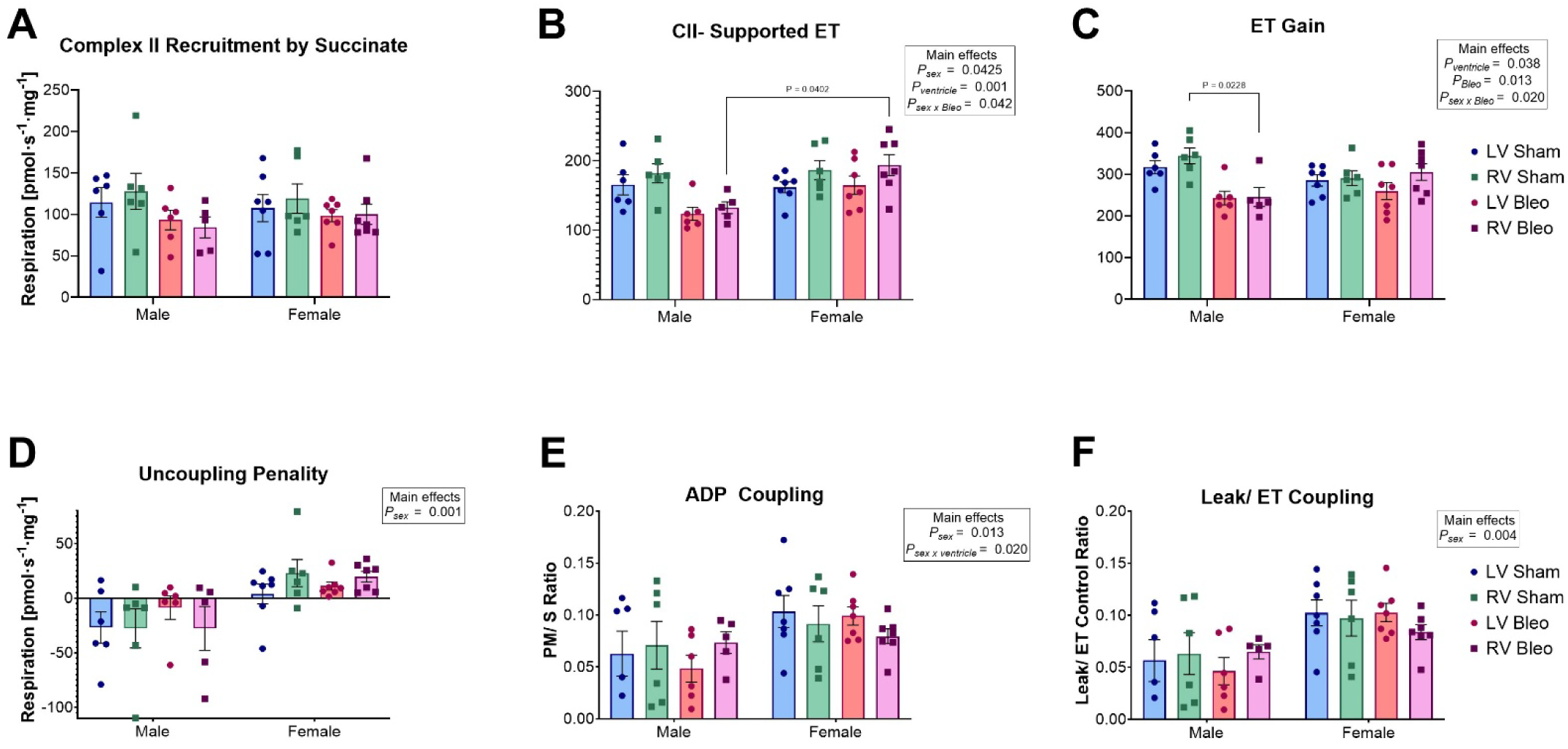
Downstream respiratory capacity and integrative control parameters in LV and RV tissue from male and female rats post-ALI are shown in (**A–F**). Complex II recruitment by succinate, calculated as the increase in respiration from NADH-linked to convergent substrate input, is shown in (**A**). CII-supported ET reflects uncoupled respiration sustained by Complex II-linked substrates (**B**), and ET gain represents the increase in respiration from OXPHOS to the uncoupled state (**C**). The uncoupling penalty indicates the absolute difference between ET capacity and OXPHOS respiration (**D**). ADP coupling describes the proportion of ET capacity achieved during OXPHOS (**E**), and the Leak/ET coupling ratio denotes the fraction of uncoupled respiration attributable to proton leak **(F)**. Bleo males showed RV reduction in CII-supported ET compared to Bleo females (**B**), and ET gain compared to Sham males (**C**). Though no significance was achieved, uncoupling penalty values exhibited a sex-dependent pattern, trending negative in males and positive in females across all conditions (**D**). No significant differences were observed in CII recruitment by succinate, ADP coupling, or Leak/ET coupling across groups. Data are expressed as mean ± SEM. Group sizes: n = 5–7 per experimental condition. Significance testing was completed using a three-way ANOVA with sex and treatment as between-subjects factors and ventricle laterality as a within-subjects factor followed by post-hoc testing using Šídák’s correction for multiple comparisons.

CII-supported ET capacity, sustained by succinate alone, showed significant main effects sex (*P_sex_* = 0.043), ventricle (*P*_ventricle_ = 0.001), and sex x treatment interaction (*P_sex x Bleo_* = 0.042); however, treatment (*P*_Bleo_ = 0.2694) did not achieve significance. Congruently, a sex difference was observed in the RV, where Bleo males exhibited lower values than Bleo females (*P* = 0.040; **Figure 4B**). This difference suggests a sex-dependent difference in CII-linked respiratory capacity in the RV following ALI.

ET gain, calculated as the increase in respiration from OXPHOS to the uncoupled state, revealed significant effects of ventricle and treatment and ventricle (*P_ventricle_* = 0.038 and *P_Bleo_* = 0.013, respectively) as well as a significant sex × treatment interaction (*P_sex x Bleo_* = 0.020). Post-hoc testing indicated a non-significant reduction of ET gain in the LV of Bleo males compared to Sham (*P* = 0.093) and a significant reduction in the RV controls (*P* = 0.023; **Figure 4C**). No differences were observed within the female groups or in the sex comparison. Although ET reserve was diminished, indices of downstream coupling control remained unaffected across experimental groups.

Uncoupling penalty, defined as the difference between uncoupled and ADP-stimulated respiration, demonstrated a significant effect of sex (*P_sex_* = 0.001). While no significant between-group differences were identified by post-hoc testing; a near-significant decrease was identified in the RV of Sham males compared to females (*P* = 0.0501; **Figure 4D**). Significant main effects of sex were observed for ADP coupling efficiency (*P_sex_* = 0.039) with a significant interaction between sex and ventricle (*P_sex x ventricle_* = 0.027; **Figure 4E**). Sex also significantly affected Leak/ET coupling ratio (*P_sex_* = 0.004). In parallel, the LV of Bleo males trended lower in the Leak/ET coupling ratio compared to Bleo females but did not achieve significance (*P* = 0.065; **Figure 4F**). Other derived metrics were largely preserved across groups. CI-mediated OXPHOS and CI vs. CII balance showed no significant differences by treatment, sex, or ventricle (**Figure S1A-B**). A significant main effect of sex was observed for the OXPHOS-ET (*P/E*) control ratio (*P_sex_* = 0.001) and trended lower in the RV of Sham males compared to females (*P* = 0.0823; **Figure S1C**). The respiratory control ratio showed only a significant interaction of sex x ventricle x treatment (*P_sex x ventricle_* _x Bleo_ = 0.049; **Figure S1D**). Though no significant changes were found following post-hoc testing, the phosphorylation (P-L) control efficiency demonstrated a significant main effect of sex (*P_sex_* = 0.039) and a sex x ventricle interaction (*P_sex x ventricle_* = 0.027; **Figure S1E**). These results indicate that while ALI impairs select elements of ET reserve and CII-linked capacity, core respiratory coupling mechanisms remain functionally intact.

### Ventricle-Specific Reduction in Mitochondria-Associated Proteins Following ALI

To investigate if mitochondrial-associated protein expression was correlated to the functional mitochondrial respiration changes, we employed Western immunoblotting (**Figure 5**). When compared to Sham, the LV of Bleo males showed no significant changes in the expression of the OXPHOS cocktail used to probe NDUFB8 (CI), SDHB (CII), UQCRC2 (CIII), MTCO1 (CIV), or ATP5A (CV; **Figure 5A**). Further probing of the ET showed no changes of NDUFS1, SDHA, or UQCRFS1 in the LV. Interestingly, BNIP3, associated with regulating mitochondrial quality control, was significantly increased in Bleo males compared to Sham (*P* = 0.007; **Figure 5B**). Conversely, the RV of Bleo males displayed significant reductions in the expression of NDUFB8 (*P* = 0.0053), UQCRC2 (*P* = 0.0008), MTCO1(*P* = 0.0054), and ATP5A (*P* = 0.046) compared to Sham males. (**Figure 5C**). In contrast, no significant changes were observed in NDUFS1, SDHA, or UQCRFS1 between Sham and Bleo in the RV; however, BNIP3 expression was markedly increased (*P* = 0.006; **Figure 5D**). These results appear to show that proteins related to components of CI, CIII, and CIV of the ET are differentially affected in each ventricle, while mitochondrial quality control in increased in both.

**Figure 5.**
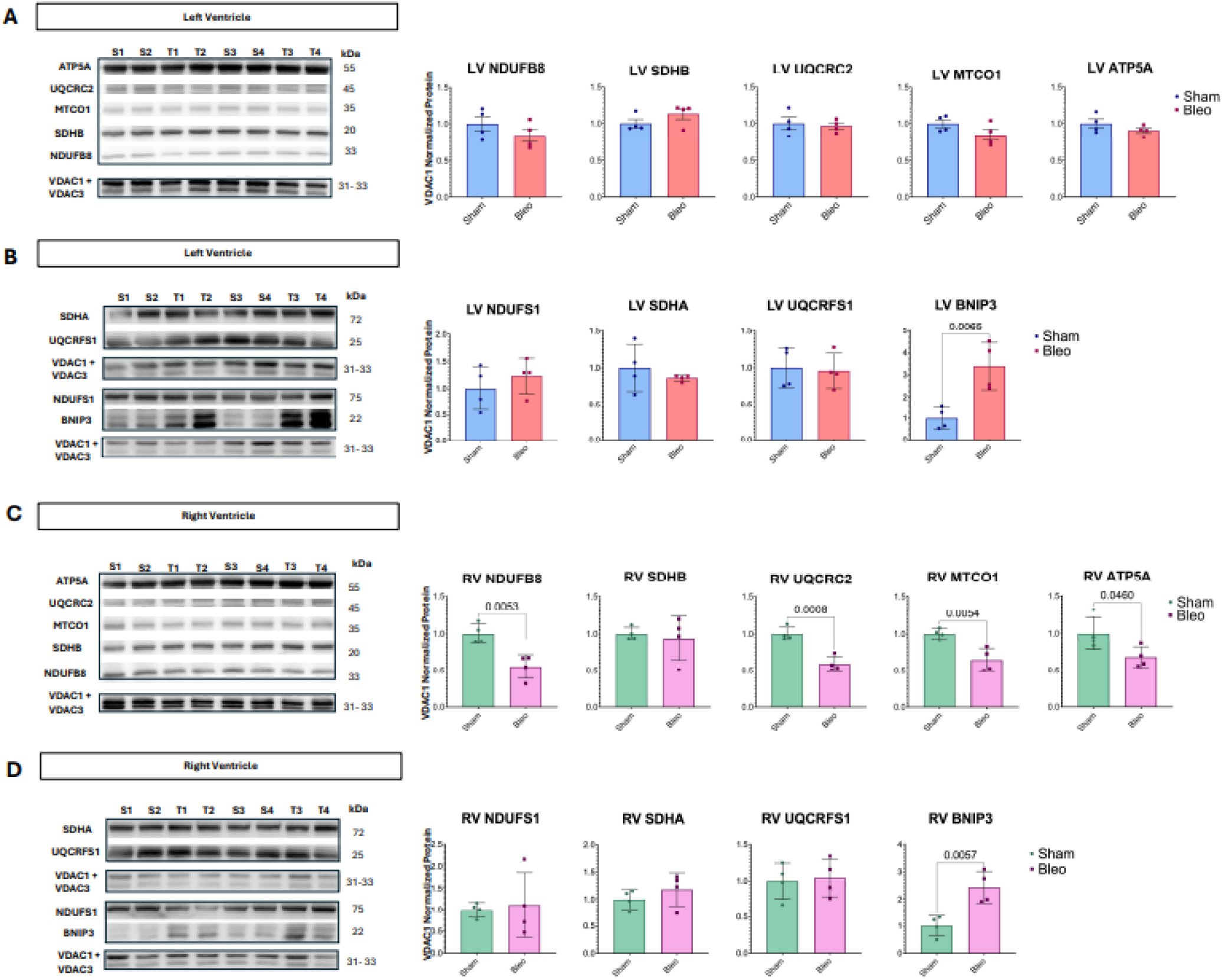
Western immunoblot images of OXPHOS proteins NDUFB8 (CI), SDHB (CII), UQCRC2 (CIII), MTCO1 (CIV), and ATP5A (CV) and quantification normalized to total VDAC1 + VDAC3 mitochondrial loading control in the LV (**A, B**) and RV (**C,D**) of Bleo male rats post-ALI. Data are expressed as mean ± SEM. Group sizes: n = 4 per experimental condition. Analysis was completed using Mann-Whitney U test.

### Pro-inflammatory Cytokine Exposure Increases Proton Leak Without Altering Global Mitochondrial Respiration

We used the Seahorse X^fe^96 Cell Mitochondrial Stress Test to elucidate the role of mitochondrial function in regulating the lung-heart axis during ALI. Mitochondrial respiratory parameters (basal respiration, non-mitochondrial oxygen consumption, proton leak (uncoupled respiration), ATP production (coupled respiration), maximal respiration, and spare respiratory capacity) were measured in H9C2 cells incubated with PRO, ANTI, and BOTH cytokine cocktails. Following a 6-hour incubation with PRO, ANTI, and BOTH treatment conditions, our results showed a slight, non-significant increase across these parameters when compared to the CTRL group (**Figure S2**). A significant increase in uncoupled respiration (proton leak) was observed in PRO (*P* = 0.022; **Figure S2D**).

After a longer, 24-hour incubation with the aforementioned treatment conditions, PRO and BOTH showed a slight increase in all parameters, including uncoupled respiration, compared with CTRL, but no pairwise differences reached significance following correction for multiple comparisons. Additionally, PRO exerted a limited main effect on uncoupled respiration (*P_PRO_* = 0.031; **Figure S3**). We suggest that pro-inflammatory cytokine cocktails initially increase uncoupled mitochondrial respiration in H9C2 cells; however, this effect was modest with prolonged exposure.

### Hypoxia Enhances Maximal Respiration and Spare Capacity Independent of Norepinephrine and Cytokine Treatment

When challenged with hypoxia for 24 hours, H9C2 cells showed a statistically significant increase in maximal respiration (*P* = 0.017; **Figure S4F**) due to an increase in spare respiratory capacity (*P* = 0.005; **Figure S4G**), but only marginal increases in non-mitochondrial oxygen consumption and basal (uncoupled and coupled) respiration.

To explore the effects of β-adrenergic receptor stimulation in H9C2 cells, we measured mitochondrial respiration following treatment with NE alone and in combination with PRO under normoxia or hypoxia for 24 hours to closely resemble physiological conditions in ALI. H9C2 cells incubated with PRO alone under both hypoxia and normoxia displayed a marginal difference compared to the control, similar to the pattern in **Figure S3**. Treatment with chemical mediators, with NE or with NE plus PRO, revealed a marginal, non-significant effect on H9C2 cells under the respective oxygen-level condition (**Figure 6**). This trend was validated in a repeat experiment under normoxic conditions (**Figure S5**).

**Figure 6.**
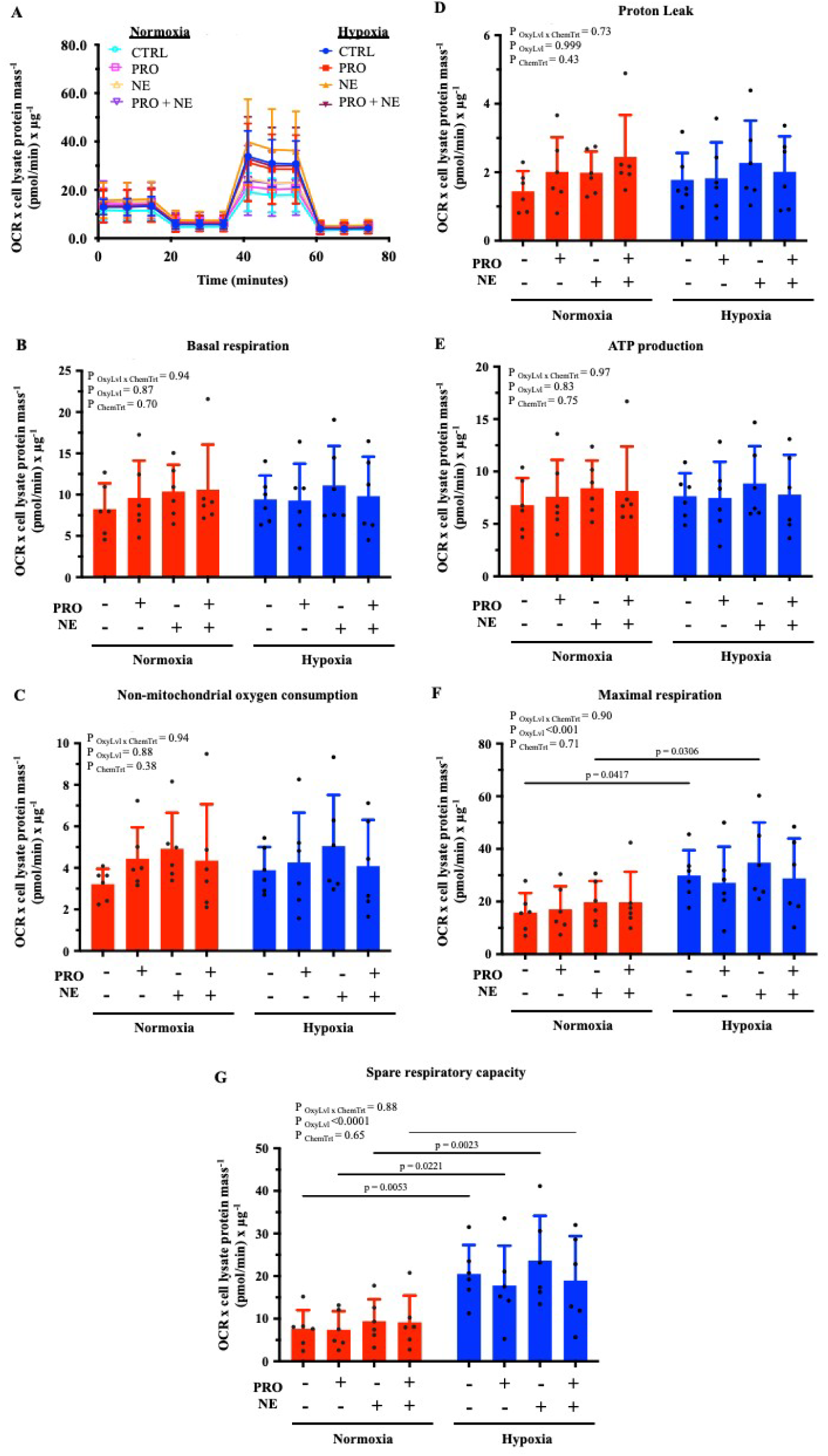
Effect of 24-hr incubation of 50µM norepinephrine (NE) and/or 100ng/ml pro-inflammatory cytokine cocktails (PRO; 100ng/mL TNFα plus 100ng/mL IL-1βplus 100ng/mL IL-6) on (**A**) mitochondrial respiration rate of H9C2, *Rattus Norvegicus* cardiomyoblast cell line at normoxia and hypoxia (1% O_2_) condition. (**B**) Basal respiration, (**C**) non-mitochondrial oxygen consumption, (**D**) proton leak, (**E**) ATP production, (**F**) maximal respiration, and (**G**) spare respiratory capacity. Data are presented as mean ± SD. Group sizes: n = 6 per experimental condition. Statistical testing was performed using two-way ANOVA, with Tukey’s multiple comparison test for post-hoc testing.

Overall, hypoxic conditions increased the maximal mitochondrial respiration in H9C2 cells, driven by a significant increase in spare respiratory capacity (**Figure 6F-G**). Indeed, maximal respiration was increased in hypoxic groups without NE or/and PRO (*P* = 0.042) and with NE alone (*P* = 0.031) compared to the respective treatments (**Figure 6F**). This corresponds with an increase of spare respiratory capacity in the hypoxia groups without NE or/and PRO (*P* = 0.005), with PRO alone (*P* = 0.022), with NE alone (*P* = 0.002), and with both PRO and NE (*P* = 0.030) relative to the respective treatments in normoxia (**Figure 6G**). Additionally, oxygen demonstrated a significant effect on the maximal respiration (*P* _OxyLvl_ = 0.0009) and spare respiratory capacity (*P* _OxyLvl_ <0.0001) in H9C2 cells regardless of the chemical mediator treatment (**Figure 6**). This implies that the hypoxic condition, rather than treatment, drives alterations in mitochondrial function.

## Discussion

This study examined ventricular mitochondrial bioenergetics during the acute phase of ALI. Our analyses revealed that the LV and RV are differentially susceptible to diminished mitochondrial function in the acute phase of ALI. Following Šídák’s corrections for multiple comparisons, PGMS*_E_* in the Mitopathways analysis showed a significant reduction in the LV of Bleo male compared to Sham. This pattern is corroborated by our Western blot data, which showed no significant alterations in ET-associated protein expression in the LV of Bleo males compared to Sham. In contrast, the RV of Bleo males exhibited impaired function across PGMS*_P_* in the Mitopathways analysis, OXPHOS capacity, ET capacity, maximal capacity, CI-supported ET, and ET gain relative to Sham males. Furthermore, Bleo males demonstrated significant reductions in protein expression of NDUFB8, UQCRC2, MTCO1, and ATP5A compared to Sham, corresponding to CI, CIII, CIV, and CV, respectively. This gives the appearance that the RV experiences markedly impaired bioenergetic function, while the LV seems to be relatively preserved. Bleo females only showed a reduction in the CI gain of the LV, compared to the RV, demonstrating a greater preservation of mitochondrial respiratory capacity compared to their male counterparts.

A sex-dependent bioenergetic response to pulmonary injury was observed. The Mitopathways metrics PGM*_P_*, PGMS*_P_*, and S*_E_*, as well as the functional indices OXPHOS capacity, OXPHOS gain, CI gain, and CII-reliant ET, appeared to be preserved in the RV of Bleo females compared to Bleo males. These mitochondrial alterations were accompanied by elevated cTnI levels in male animals, suggesting early cardiac stress following ALI.

Lung diseases, including ARDS, have long been associated with cardiovascular complications^15^. Cardiac stress may arise not only from hemodynamic strain but also from circulating inflammatory mediators and oxidative stress. Elevated cardiac troponin (hsTn) and cTnI, used to detect myocardial injury, have been observed in over 90% of patients with ARDS^16–18^. We measured cTnI seven days post-ALI and found elevated levels in male Bleo rats compared to both Sham and female Bleo groups (**Figure 1**). These findings indicate that cardiac stress develops during the acute phase of lung injury.

Although experimental models such as hypoxia, sepsis, and ischemia-reperfusion injury differ from ALI, they provide important mechanistic insight into cardiopulmonary stress and mitochondrial dysfunction. Hypoxemia is a defining feature of ALI, arising from impaired pulmonary gas exchange^19^. Both sepsis and ischemia-reperfusion injury are associated with impaired cardiac mitochondrial respiration and coupling efficiency, frequently characterized by decreased OXPHOS capacity, increased mitochondrial oxidative stress, and cytochrome c release into the cytoplasm^20,21^.

We evaluated oxygen consumption in permeabilized cardiac muscle fibers from the LV and RV of rats to investigate the significance of metabolic and mitochondrial function in ALI-induced cardiac injury. Mitochondrial respiration in the LV appeared to show modest changes following Bleo. Mitopathways analysis revealed reduced PGMS*_E_* capacity in Bleo males (**Figure 2B**); however, Leak respiration, OXPHOS capacity, ET capacity, and CIV activity were relatively preserved. Bleo females likewise demonstrated preserved respiratory function across all measured parameters (**Figure 2D-G**). To our knowledge, studies assessing HRR in cardiac tissue in ALI models remain inchoate, but patterns of relatively preserved LV mitochondrial respiration have been reported in other cardiopulmonary stress models^22–24^. For example, Horscroft et al.^22^ observed alterations in baseline Leak respiration following sustained hypoxic exposure, while other respiratory parameters in the LV remained largely unchanged in male rats. Studies examining other forms of cardiac injury have reported pronounced mitochondrial impairment in the LV. For example, Szibor et al.^23^ demonstrated significant reductions in CI-linked OXPHOS and CIV activity following ischemia-reperfusion injury in the LV of male mice. Although this model represents a direct cardiac insult rather than bleomycin-induced ALI, these findings highlight that mitochondrial respiratory responses in the LV may vary with the underlying pathological stimulus.

In our study, several respiratory control indices showed non-significant downward trends among Bleo males, including OXPHOS gain, CI gain, ET gain, and CI- and II-supported respiration. These same respiratory control indices were analyzed in Bleo females as well, which did not exhibit measurable changes. We further investigated specific proteins involved in CI and II respiration. Changes in protein expression of NDUFB8, the gene-coding protein associated with the structural integrity and assembly of CI, nor NDUFS1, which directly facilitates the primary catalytic activity of CI, were not observed in the LV of Bleo males relative to Sham (**Figures 5A-B**). Concomitant downstream functional mitochondrial respiration metrics CII recruitment by succinate under OXPHOS, CII-supported ET capacity, ET gain, uncoupling penalty, ADP coupling, and Leak/ET coupling ratio in the LV revealed no significant changes in Bleo males compared to Sham (**Figure 4**). Though no significance was observed in CII-recruitment by succinate under OXPHOS, the overall magnitude of convergent electron flow through Complexes I and II increased OXPHOS capacity relative to CI-supported respiration alone in both sexes (**Figure 4A**). Additionally, there were no significant differences seen in CII complex proteins SDHA and SDHB. Sequentially, UQCRC2 and UQCRSF1, both integral to CIII, showed no significant differences in protein expression in the LV of Bleo males compared to Sham. By extension, proteins involved in CIV and CV, MTCO1 and ATP5A, respectfully, were analyzed and found to not have significant changes in protein expression in our model (**Figures 5A-B**).

More severe mitochondrial dysfunction has also been reported in systemic inflammatory injury; for example, Doerrier et al.^24^ demonstrated marked reductions across OXPHOS and ET capacities in the LV of male mice during sepsis. Collectively, these observations suggest that the extent of LV mitochondrial respiratory impairment varies according to the nature and severity of the underlying pathological stimulus.

The RV is particularly susceptible to lung-derived injury during ARDS, a condition associated with increased mortality, as pulmonary vascular dysfunction and elevated afterload impose sustained metabolic stress on the myocardium^25,26^. Mitochondrial metabolic remodeling and reductions in oxidative phosphorylation capacity have been linked to impaired energetic reserve and contractile dysfunction in the RV during pulmonary hypertension and pressure overload^27,28^.

Alongside changes in mitochondrial respiration observed in the LV, bioenergetic impairment was also observed in the RV. Absolute respiratory flux measurements were markedly suppressed, indicating reduced mitochondrial oxidative capacity in Bleo male rats (**Figure 2**). Mitopathways analysis further demonstrated reduced respiratory flux in PGMS*_P_* and near significant reductions of PGMS*_E_* and S*_E_* (**Figure 2C**). Reductions in substrate-supported OXPHOS, ET capacity, and maximal respiratory capacity, together with preserved Leak and CIV activity in the male RV, suggest a loss of mitochondrial reserve that extends beyond dysfunction within a single respiratory pathway (**Figure 2D-G**).

Although glutamate-driven CI respiration exhibited only a downward trend, the RV of Bleo males showed reduced ET gain and diminished CI-supported ET capacity, indicating impairment at the level of CI-linked electron transport (**Figure 3**). We observed reduced protein expression of NDUFB8, in RV of Bleo males compared to Sham (**Figure 5C**). Interestingly, there was no change in the expression of NDUFS1 (**Figure 5D**).

This pattern is recapitulated in prior studies, where similar mitochondrial respiratory alterations involving CI have been described in other forms of right ventricular dysfunction^29,30^. For example, Hwang et al.^29^ reported reductions in OXPHOS and uncoupled respiration in human samples from patients with severe RV hypertrophy and failure, while glutamate-driven CI respiration remained unchanged, and succinate-supported CII respiration was preserved. Similarly, Kumari et al.^30^ observed reduced CI-stimulated respiration with preserved CII- and CIV-mediated respiration in the RV at postnatal day 21 in a postnatal hyperoxia rat model. Although these studies were conducted in different models of cardiopulmonary stress, they highlight selective remodeling of mitochondrial respiratory pathways in the RV, particularly involving CI-linked respiration.

The selective reduction in respiration supported by NADH-linked substrates, along with diminished CI gain and CI-supported ET, suggests that ALI preferentially impairs CI-mediated control of mitochondrial respiration in RV. In contrast, CII-linked respiration and downstream coupling efficiencies appear largely preserved, indicating that the primary site of respiratory disruption lies upstream, at or before CI entry. This is further validated by the preservation of convergent electron flow through Complexes I and II via CII-recruitment by succinate under OXPHOS in both sexes (**Figure 4A**), which demonstrated an increase of OXPHOS capacity relative to CI-supported respiration alone. Indeed, we saw no changes in expression of CII-associated proteins SHDA, nor SDHB, which transfer electrons to coenzyme Q in the ET (**Figures 5D-E**). However, reductions in protein expression of downstream ET complexes III, IV, and V were found. The expression of UQCRC2 was found to be reduced in the RV of Bleo males compared to Sham; however, UQCRFS1, which directly interacts with UQCRC2 and CIV, showed no significant changes. Perhaps the limited reduction in functional metrics shown in HRR could, in part, be attributed to only one of these proteins showing altered expression. Additionally, we demonstrated reductions in both MTCO1, the catalytic core of CIV, and ATP5A, the catalytic subunit of CV. Moreover, we determined the protein expression of BNIP3, which is associated with mitochondrial quality control and mitophagy, was elevated in Bleo males compared to Sham (**Figure 5D**). Potential issues and disruptions to mitochondrial quality-control pathways, such as mitophagy and ferroptosis, can worsen metabolic remodeling and contribute to lung-derived inflammatory damage during ALI^31^. Taken together, these results show that Bleo causes asymmetric myocardial injury and ventricular dichotomy in mitochondrial respiratory capacity and control, with the RV emerging as a site of heightened vulnerability.

Sex-specific differences in cardiopulmonary adaptation may contribute to the divergent mitochondrial phenotypes observed in this study. In conditions that increase pulmonary vascular load, the sex hormone estrogen was found to protect RV, leading females to demonstrate improved function and metabolic adaptation compared with males^32,33^. In cardiac inflammatory models, females demonstrate greater enrichment of mitochondrial metabolic pathways, whereas males show increased inflammatory signaling and reduced mitochondrial respiratory gene expression^34^.

Sex-dependent differences in mitochondrial respiration were evident following lung injury. The RV of female Bleo rats demonstrated preserved absolute respiratory flux and a divergent ventricular response of increased CI gain in the RV relative to the LV (**Figure 3**). No changes were observed in Leak respiration or CIV activity, suggesting that the mitochondrial membrane proton gradient and terminal electron transport remain intact. This is consistent with prior work demonstrating enhanced mitochondrial resilience in females under cardiopulmonary stress, including improved mitochondrial regulation and adaptive responses in pulmonary hypertension models^35^.

In the Mitopathways comparison of respiratory flux, the RV of Bleo males exhibited reduced respiratory flux in PGM*_P_* and PGMS*_P_* capacity and lower S*_E_* capacity than females, indicating diminished NADH- and succinate-linked oxidative capacity (**Figure 2C**). Sex differences in the LV appeared to be modest, with Bleo males showing reduced PM*_L_* compared with females, while other respiratory states remained largely unchanged.

Respiratory control metrics further emphasized sex-dependent RV alterations. Compared with Bleo males, Bleo females demonstrated greater OXPHOS capacity and higher CI gain, whereas males showed reduced OXPHOS gain (**Figure 3**) and lower CII-supported ET capacity (**Figure 4**). Glutamate-driven CI respiration trended lower in males but did not reach significance. We found no significant changes in the LV; however, Bleo males trended toward lower Leak respiration and reduced ADP coupling. The uncoupling penalty showed no group differences but trended negatively in males and positively in females, while the Leak/ET ratio trended higher in females. Together, these findings indicate a ventricle-specific, sex-dependent mitochondrial phenotype, with males demonstrating reduced oxidative capacity and respiratory reserve in the RV. At the same time, females maintain greater bioenergetic stability under pulmonary injury^36^.

Despite limited studies using HRR to compare sex differences in ALI models, several established hypotheses may contribute to the observed protective effect in females. Di Florio et al. shows that in viral myocarditis female hearts exhibit upregulation of mitochondrial biogenesis and energy metabolism genes, while enriched inflammatory pathways in males reflected worse electron transport^34^. Mechanistically, the sex hormone estrogen has been linked to the regulation of mitochondrial biogenesis and oxidative phosphorylation, including increased expression of key mitochondrial regulatory genes such as PGC-1α, nuclear respiratory factor-1 (NRF1), and estrogen-related receptor-α (ERRα)^37,38^. In pulmonary hypertension, estrogen protects RV function and mitigates metabolic remodeling in rats^33^. Specifically, estradiol has been shown to preserve RV function and maintain mitochondrial metabolic capacity in models of pulmonary vascular disease^39^. Furthermore, 17β-estradiol reduces the remodeling of glutamatergic vagal afferent neurons following myocardial infarction (MI) in female mice, leading to decreased mitochondrial dysfunction and oxidative stress compared with males^40^.

Although the specific effects of sex hormones in ALI models remain to be investigated, we have exhibited concurrent evidence of a female-protective mechanism in response to ALI. This observation is consistent with recent clinical studies of SARS-CoV-2 infection, demonstrating that males experience greater disease severity and higher mortality than females despite similar infection rates, suggesting enhanced resilience to severe viral lung injury in females^41–43^.

Moreover, our findings of diminished mitochondrial function in the LV and RV during the acute phase of ALI align with emerging evidence that severe lung injury can induce systemic mitochondrial dysfunction across multiple organs. Previous studies of SARS-CoV-2 analyzed autopsy tissue of the heart, liver, kidney, and lymph nodes for mitochondrial bioenergetic gene expression, revealing downregulation in OXPHOS mRNAs, with the most downregulation occurring in the heart^44^. Widespread mitochondrial dysfunction has been identified as a unifying mechanism linking oxidative stress and cardiac injury in COVID-19 patients, characterized by attenuated oxidative phosphorylation, disrupted mitochondrial ultrastructure in both cardiac and skeletal muscle, and a shift toward glycolysis in cardiac tissue, in part via upregulated HIF-α1^4,45–47^. Although our *ex vivo* HRR study validates the contribution of mitochondrial impairment to lung-heart crosstalk, the mechanistic pathway involving molecular mediators remains to be elucidated.

The role of macrophage phenotypes in myofibroblast differentiation has been identified by utilizing single-cell RNA sequencing of cardiac immune cells. Pro-inflammatory macrophages rely predominantly on glycolysis, whereas anti-inflammatory macrophages favor fatty acid oxidation and oxidative phosphorylation^48^. Further, high serum levels of proinflammatory cytokines, particularly IL-6 and TNF-α, contribute to systemic inflammation and are strong predictors of patients’ survival, serving as significant predictors of disease severity and death even after adjusting for clinical status, inflammation and hypoxia, demographics, and comorbidities^44,45,49^. Catecholamines, such as NE, have been shown to participate in the development of ARDS through AR activation^50^. In COVID-19 patients, NE and IL-6 are significantly correlated with the highest risk of mortality^51,52^.

To further investigate these effects, we analyzed mitochondrial respiration using Seahorse X^fe^96 Cell Mitochondrial Stress Test in the H9C2 *Rattus Norvegicus* cardiomyoblast cell line after short and long exposure to PRO, ANTI, and BOTH cytokine cocktails. This was based on the dynamics of pro- and anti-inflammatory cytokines observed to be associated with ALI pathogenesis^53^. A surge of pro-inflammatory cytokines, such as IL-6, IL-1β, and TNFα, was observed in plasma during ALI^54–56^. This systemic cytokine storm could expose other organs, including cardiac tissues, to high concentrations of inflammatory cytokines^57^. An elevated inflammatory response is critical because crosstalk between cytokines and mitochondrial function has been reported in injured tissues and disease states ^58–61^. Subsequently, we hypothesized that inflammatory cytokines exert differential effects on mitochondrial respiration in H9C2 cells following exposure to high-dose cytokine cocktails.

We demonstrated an increase in uncoupled respiration following a 6-hour incubation of PRO compared to the CTRL (**Figure S2D**); however, no significant differences were found across the remaining mitochondrial respiratory parameters of basal respiration, non-mitochondrial oxygen consumption, coupled respiration, maximal respiration, and spare respiratory capacity (**Figures S2-S3**). Additionally, a modest effect of PRO on uncoupled respiration was observed after 24-hour incubation (**Figure S3D**). These results appear to contrast with reduced Leak respiration demonstrated by overall main effect of sex in both ventricles (**Figure 2D**) but aligns with the significant increase of PM*_L_* in the LV, and trend towards significance in the RV of Bleo females compared to males in the Mitopathways analysis (**Figure 2B-C**).

To our knowledge, there are no current studies that have used the Mitochondrial Stress Test to examine H9C2 cells under these conditions. In a similar study, Szczesnowski et al. (2025) illustrated significant increases in proton leak and basal respiration, and a significant decrease in coupling efficiency in the H9C2 cell line following exposure to the inflammatory mediator lipopolysaccharide (LPS) at a lower concentration for 24 hours, but the effect diminished at a high concentration over a longer exposure period^62,63^. Although proton leak has been studied in the heart, the expression of uncoupling proteins has led to controversial findings controversial regarding its beneficial or detrimental effects^64,65^. This underscores the need for further investigation to determine whether our approach to heightening proton leak in cardiomyoblasts by a shock of pro-inflammatory cytokines has a therapeutic or deleterious impact.

During ALI, lung tissue inflammation and leukocyte infiltration increase alveolar-epithelial barrier permeability, followed by pulmonary infiltration and edema^66,67^. This results in hypoventilation, pulmonary shunting, and hypoxia in other organs and tissues, including the cardiac tissue^68^. Cardiac hypoxia modulates metabolic adaptation mechanisms, including regulation of mitochondrial function^69–71^. To investigate this, we compared mitochondrial respiration in the H9C2 cell line after incubation under normoxia and hypoxia for 24 hours.

Under hypoxic conditions, we observed significant increases in maximal respiration, spare respiratory capacity, as well as marginal changes in non-mitochondrial oxygen consumption and uncoupled and coupled basal respiration (**Figure S4**). A similar trend was observed under these conditions, wherein maximal respiration and spare respiratory capacity were increased, independent of the chemical mediator treatment (**Figure 6**). This phenomenon was not reflected in the *ex vivo* HRR experiment, in which maximal respiration was decreased in the RV of Bleo males compared to Sham males (**Figure 2G**). Nevertheless, a recent study has shown a similar phenomenon by investigating Human c-Kit+ cardiac progenitor cells (hCPCs) isolated from the LV and maintained under permanent hypoxia (hCPC-1%). Significantly higher basal, ATP-coupled, and maximal uncoupled OCRs were observed compared to cells maintained under normoxia ^72^.

Lung tissue injury is accompanied by hyperactivation of the sympathetic nervous system with the secretion of catecholamines, across the brain-neural-lung axis ^73–75^, and is associated with elevated serum NE^3^. Additionally, according to Demchenko et al., the sympathetic outflow, mediated by the central nervous system during lung injury could lead to LV dysfunction^76^. Cardiac sympathetic nerve activity with NE spillover is strongly related to cardiac injury and failure severity, and β-adrenergic receptor stimulation contributes to the regulation of mitochondrial biogenesis and function ^70,77–80^. Therefore, we investigated the implications of β-AR stimulation by treating H9C2 cells with NE alone and in combination with PRO under normoxia or hypoxia for 24 hours.

We observed no significant changes in mitochondrial respiration after treatment with NE or in combination with PRO under both normoxic and hypoxic conditions compared to the respective control groups (**Figures 6, S5**). This contrasts with our *ex vivo* HRR results, which showed diminished mitochondrial function in the LV and RV post-ALI. Nonetheless, studies utilizing different approaches to measure mitochondrial function showed that NE induces ROS-dependent hypertrophic and apoptotic responses in H9C2 cells and *ex vivo* cardiac myocytes^81–83^. Further, NE may directly impact mitochondrial bioenergetics by altering electron transport chain efficiency and promoting proton leak across the inner mitochondrial membrane^84^. Moreover, long-term NE (in days) exposure, in conjunction with LPS stimulation, exacerbated alterations in ROS levels, mitochondrial shrinkage, and ferroptosis observed in H9C2 cells and *in vivo*, rather than at early stages^85^. This suggests that the experimental limitation of our study, using a 24-hour NE incubation time, may require a longer exposure window for adrenergic stress to exert its effects on H9C2 cells.

Given that mitochondrial respiration and redox balance are tightly coupled with the cellular microenvironment, oxygen availability in standard cell culture practice introduces a variable that may influence baseline metabolic and inflammatory states^86^. Manipulation of both O₂ and CO₂ levels, particularly under hypoxic conditions, could modulate pro-inflammatory cytokine expression.

Our assessment of cardiac mitochondrial function focused on the acute post-injury phase, as previous work from our laboratory identified spontaneous arrhythmias at this time point, which resolve four weeks after injury^9^. HRR in permeabilized cardiac fibers provides detailed measurements of substrate-specific respiratory capacity and bioenergetic indices, complemented by the Seahorse X^fe^96 Cell Mitochondrial Stress Test to provide mechanistic insight into respiratory responses. It should be noted that mitochondrial ultrastructure, dynamics, and ROS production were not directly assessed by these approaches.

We acknowledge that our current data does not confirm detrimental effect of pro-inflammatory milieu or sympathetic stimulation on mitochondrial respiratory function. This may be, in part, because the H9C2 cell line remains immobile, rather than differentiated. Furthermore, due to the maximal treatment exposure time measured in our experiments of 24 hours, later decompensation following prolonged exposure cannot be excluded. In addition, despite increased circulating cTnI levels in male animals, we did not perform direct molecular analyses to evaluate cardiomyocyte injury or fibrosis within the myocardium. Together, these limitations indicate that our findings define an early metabolic profile of cardiac vulnerability following ALI and underscore the need for future studies to further characterize mitochondrial remodeling processes and structural cardiac injury throughout disease progression.

## Acknowledgement

We thank the Bioassay Core of the Center for Heart and Vascular Research at UNMC for providing access to and technical support for the hypoxia chamber. We gratefully acknowledge Dr. Jocelyn Plowman for her excellent editorial assistance. We also thank the Seahorse Core of the Department of Neurological Sciences at UNMC for providing access to the XFe96 Flux Analyzer and for technical support.

## Disclosure of Artificial Intelligence (AI) Assistance

The authors did not use generative AI or AI-assisted technologies in the development of this manuscript.

