## Supplement Figures for "Cardiac Mitochondrial Dysfunction Following Bleomycin-Induced Acute Lung Injury in Rats"

**Running Title: Cardiac Mitochondria Dysfunction post ALI**

### **Address Correspondence To:**

Han-Jun Wang M.D.,

Department of Anesthesiology

University of Nebraska Medical Center, Omaha, NE 68198, USA

**Figure S1:**

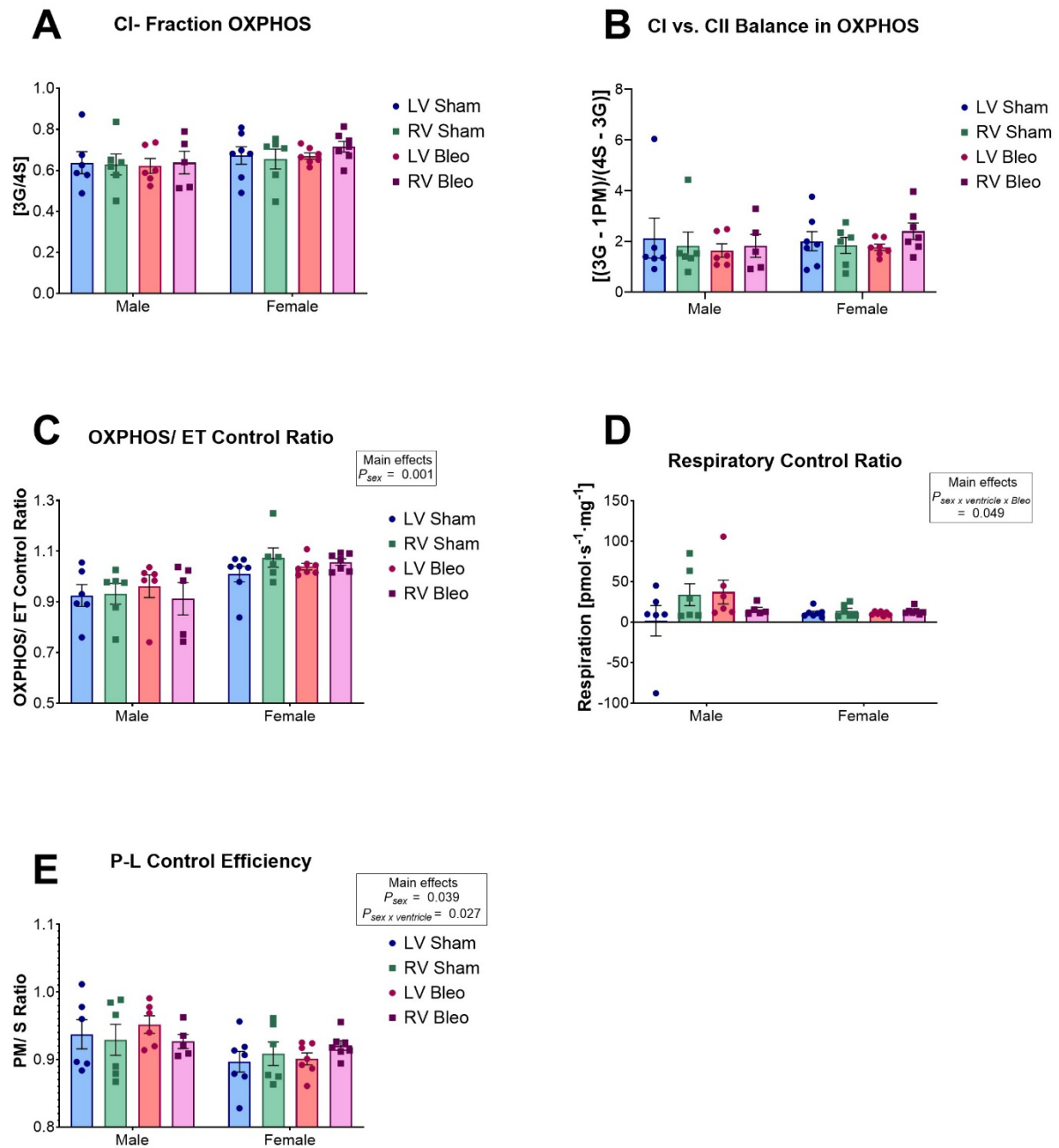

**Figure S1.** Derived mitochondrial respiratory control parameters in LV and RV tissue from male and female rats following ALI. Derived mitochondrial respiratory metrics calculated from HRR

in male and female rats post- ALI are shown in **(A–E)**. The fractional contribution of Complex I–supported respiration to total OXPHOS capacity (CI fraction OXPHOS) is shown in **(A)**. The balance between Complex I– and Complex II–supported respiration during OXPHOS is presented in **(B)**. The OXPHOS/ET control ratio, reflecting the proportion of electron transport system (ETS) capacity utilized during phosphorylating respiration, is shown in **(C)**. Respiratory control ratio (RCR), representing the ratio of OXPHOS to leak respiration, is shown in **(D)**. Phosphorylation control efficiency (P–L control efficiency), indicating the efficiency of ADP-stimulated respiration relative to proton leak, is shown in **(E)**. No significant differences were observed across treatment, sex, or ventricle for any derived metric. Data are expressed as mean ± SEM with individual data points. Group sizes: $n = 5–7$ per experimental condition. Statistical analysis was performed using three-way ANOVA with sex, treatment, and ventricle as factors, including matched LV and RV samples from individual animals, followed by Šídák's multiple comparisons test.

Figure S2

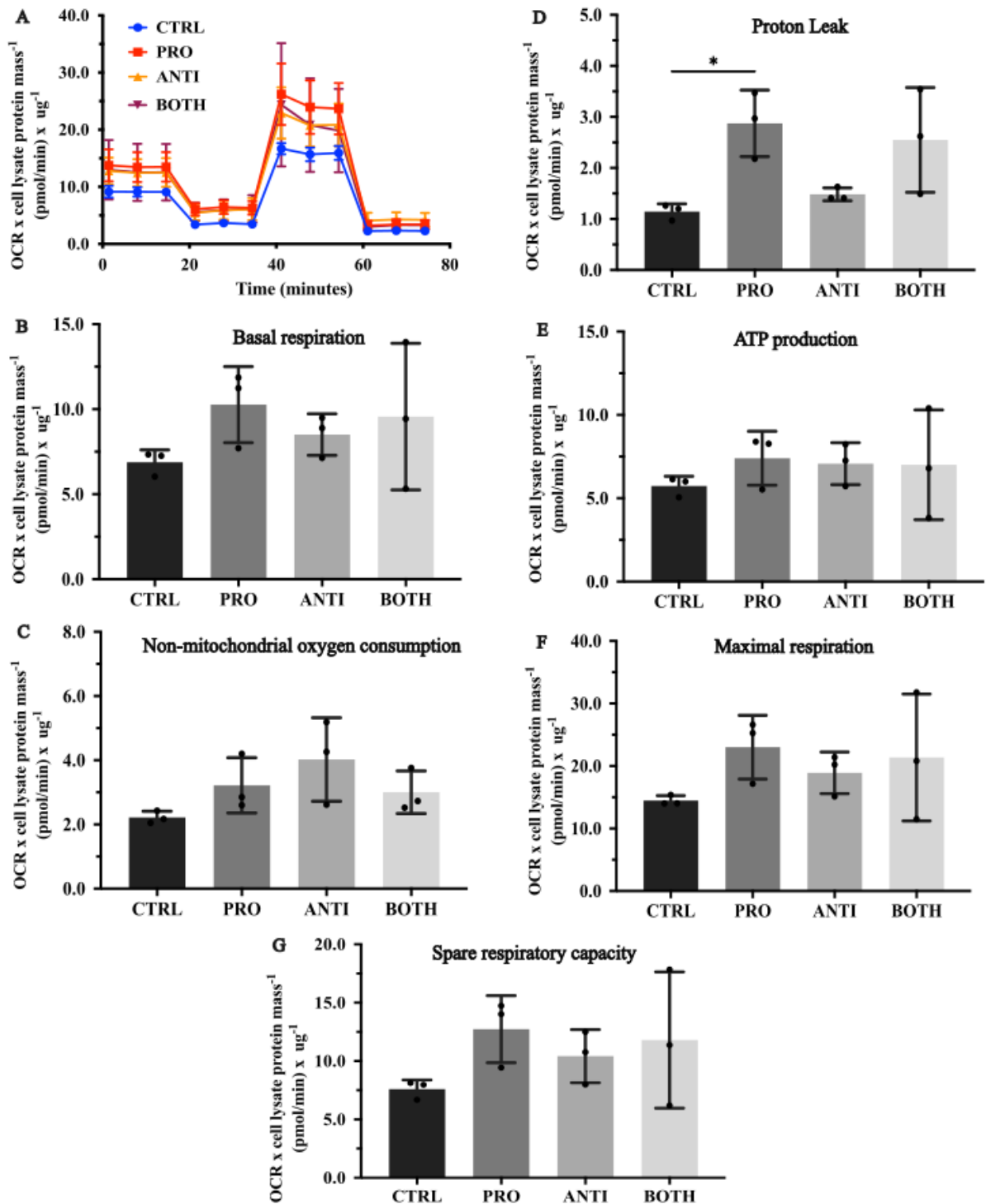

**Figure S2.** Effect of 6-hr incubation of 100ng/ml inflammatory cytokine cocktails on mitochondrial respiration rate of H9C2, *Rattus Norvegicus* cardiomyoblast cell line (A). The cells were treated with pro-inflammatory cytokine cocktails (PRO) at a final concentration of 100 ng/ml each of interleukin (IL)-6, IL-1β, and tumor necrosis factor (TNF)-α, anti-inflammatory cytokine cocktails (ANTI) at a final concentration of 100 ng/ml each of IL-10, and IL-4, or a mixture of PRO and ANTI (BOTH) for 6 or 24 hours before measurement of mitochondrial respiration rate. (B) Basal respiration, (C) non-mitochondrial oxygen consumption, (D) proton leak, (E) ATP production, (F) maximal respiration, and (G) spare respiratory capacity were quantified. Group sizes: n = 3 per experimental condition. Statistical analysis was performed using one-way ANOVA with post-hoc testing performed using Dunnett's method for multiple comparisons. Data are presented as mean ± SD. PRO, or ANTI, or BOTH, compared against CTRL: \*$P$<0.05.

Figure S3

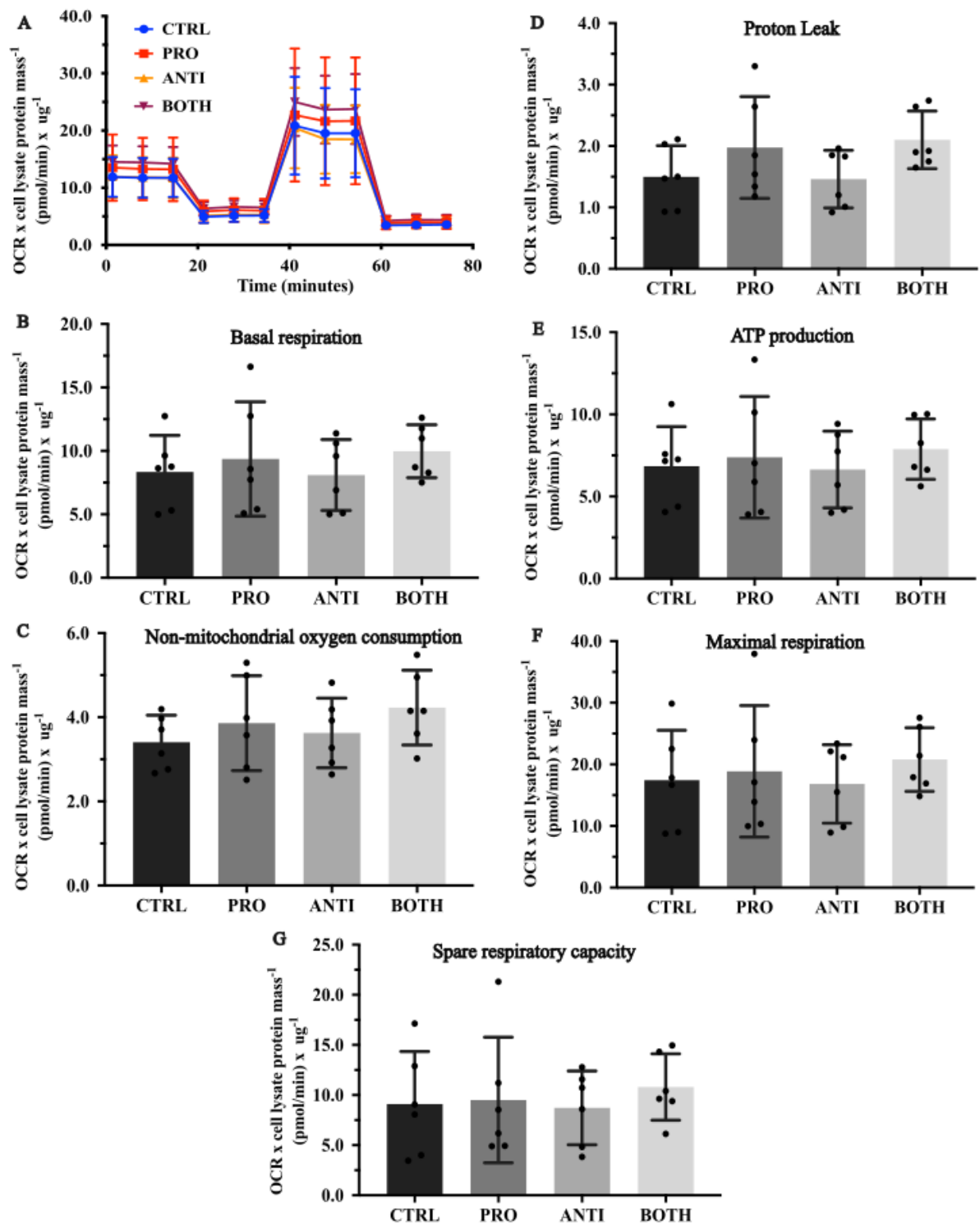

**Figure S3.** Effect of 24-hr incubation of 100ng/ml inflammatory cytokine cocktails on (A) mitochondrial respiration rate of H9C2, *Rattus Norvegicus* cardiomyoblast cell line. (B) Basal respiration, (C) non-mitochondrial oxygen consumption, (D) proton leak, (E) ATP production, (F) maximal respiration, and (G) spare respiratory capacity were quantified. Group sizes: n = 6 per experimental condition. Statistical analysis was performed using two-way ANOVA with post-hoc testing using Tukey's correction for multiple comparisons. Data are presented as mean ± SD. PRO or ANTI or BOTH, compared against CTRL: NS $P > 0.05$; Main effect and Interaction: $P_{\text{PRO}} \times P_{\text{ANTI}} > 0.05$; $*P_{\text{PRO}} < 0.05$; $P_{\text{ANTI}} > 0.05$.

**Figure S4**

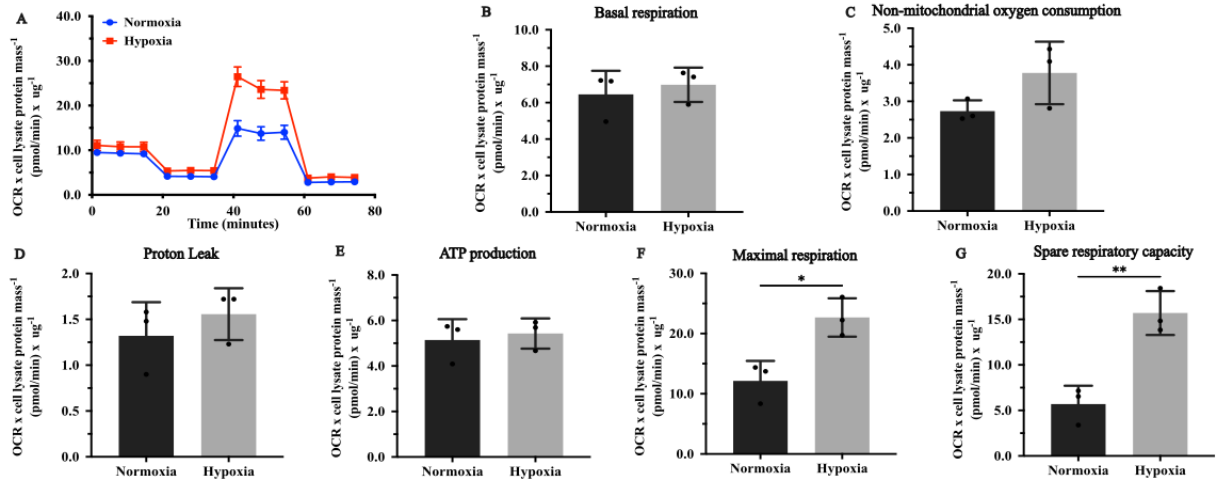

**Figure S4.** Effect of normoxia and hypoxia on (A) mitochondrial respiration rate of H9C2, *Rattus Norvegicus* cardiomyoblast cell line. (B) Basal respiration, (C) non-mitochondrial oxygen consumption, (D) proton leak, (E) ATP production, (F) maximal respiration, and (G) spare respiratory capacity were quantified. Group sizes: $n = 3$ per experimental condition. Statistical analysis was performed using an unpaired t-test. Data are presented as mean ± SD. Normoxia vs. Hypoxia: \*$P < 0.05$, \*\*$P < 0.01$.

Figure S5

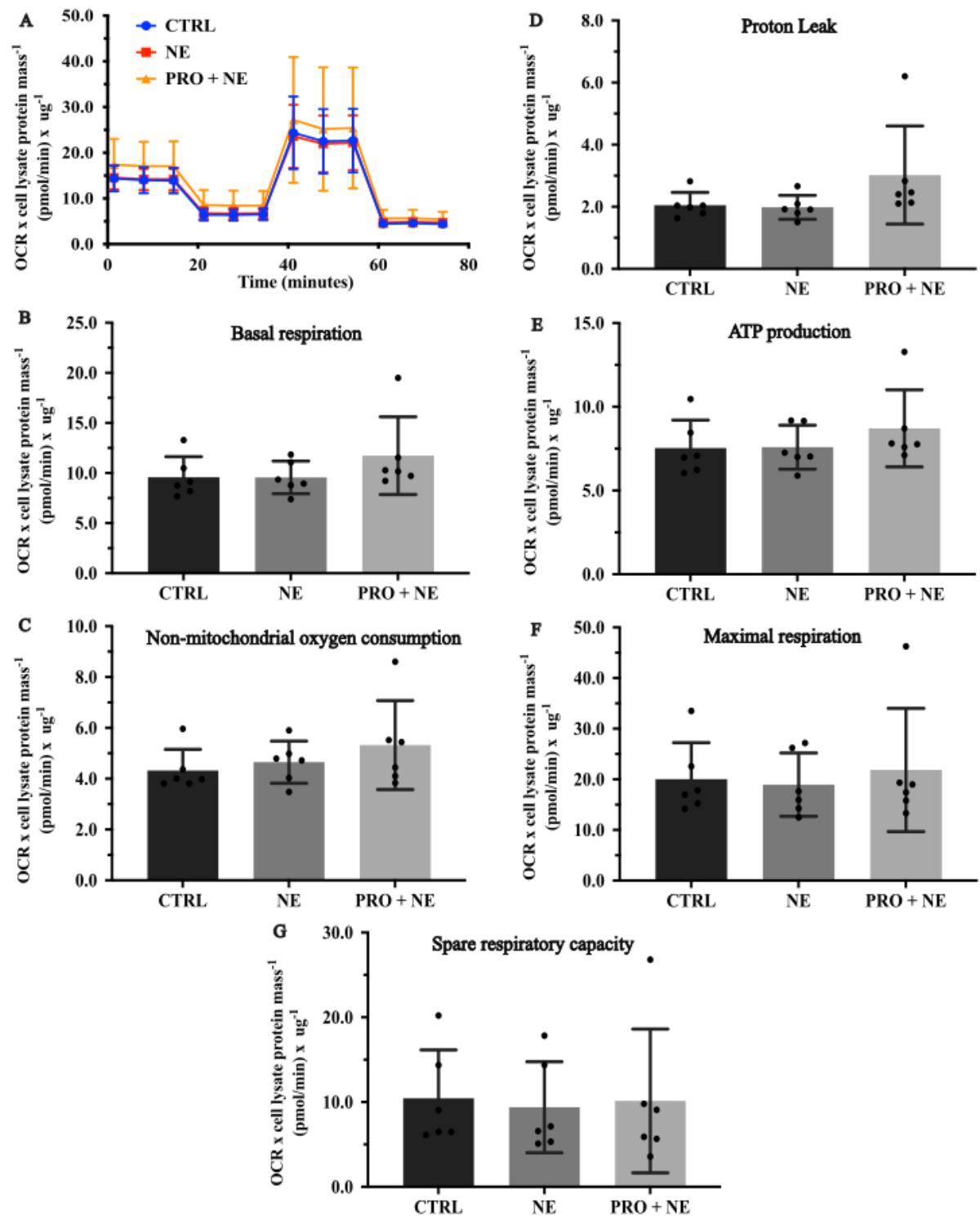

**Figure S5.** Effect of 24-hr incubation of 50μM NE and/or 100ng/ml PRO on (A) mitochondrial respiration rate of H9C2, *Rattus Norvegicus* cardiomyoblast cell line. (B) Basal respiration, (C) non-mitochondrial oxygen consumption, (D) proton leak, (E) ATP production, (F) maximal respiration, and (G) spare respiratory capacity were quantified. Group sizes: n = 6 per experimental condition. Statistical analysis was performed using one-way ANOVA with post-hoc testing using Dunnett's method for multiple comparisons. Data are presented as mean ± SD. NE or PRO, compared against CTRL: NS $P > 0.05$.
